# A test for linkage disequilibrium in structured populations

**DOI:** 10.64898/2026.10.18.759967

**Authors:** Alvin J. Zhang, John D. Storey

## Abstract

Quantifying patterns of dependence between distinct loci, known as linkage disequilibrium (LD), is vital for identifying locus–trait associations and studying evolution. LD is well understood when individuals are sampled from a population where Hardy-Weinberg equilibrium (HWE) holds at each locus. However, population structure is nearly ubiquitous in human populations, leading to significant deviations from HWE and inducing correlations between loci that mimic LD. These confounding signals represent an ongoing challenge when characterizing LD for individuals sampled from large human datasets. To address this, we develop a hypothesis test for a quantity we call *structural LD*, defined as LD after accounting for population structure. Our test accommodates arbitrarily complex structure using individual-specific allele frequencies, which are possibly distinct for every SNP–individual pair. This formalization is highly general and encompasses a wide range of established models, including admixture. We validate the performance of our proposed method on both simulated genotypes and existing human studies, including data characterized by highly complex structure and recent admixture. Our analyses of human studies indicates that most long-range LD in human populations can be accounted for by well-fit models of population structure.

## 1 Introduction

A fundamental problem in modern genetics lies in understanding and quantifying patterns of dependence across the genome in populations, known as linkage disequilibrium (LD). Patterns of LD are vital to the identification of locus–trait associations [1] and the construction of polygenic scores [2], operating via marker loci that are correlated with causal variants. LD is also indispensable for the study of evolutionary and demographic history, representing a key signature of selection and other population-level processes [3]. To this end, significant efforts have been made to produce accurate genome-wide profiles of LD in humans [4].

The process of quantifying LD is well understood when the individuals under study are sampled from a randomly mating and well-mixed population, in which each locus is under Hardy-Weinberg equilibrium (HWE) [5]. This essentially means that each locus is parameterized by a single allele frequency shared across all individuals, with expected genotype proportions determined by this frequency. Population structure gives rise to systematic deviations from HWE and can confound other sources of LD. Structure arises from a plethora of factors, including geographic subdivision, genetic drift, and finite population sizes; it can be understood intuitively as variation in allele frequencies across individuals or subsets of a population [6,7]. Because structure is nearly ubiquitous in human populations across the world [8], characterizing LD in population-sampled individuals remains an important challenge.

Population structure is well known to induce correlations between loci that mimic LD [3,9]. This is a phenomenon variously described as *spurious LD* or the *Wahlund effect at two loci* [10] and can appear even for independently assorting loci [11]. Importantly, correlations corresponding to spurious LD can confound those arising from tight genetic linkage, leading to challenges in downstream analyses (e.g., association studies) relying on the latter. When population structure consists of discrete, non-overlapping subpopulations, such as when ancestry labels are available, existing methods for characterizing LD perform well by evaluating each subpopulation separately [4,12]. A more general scenario arises when individuals cannot be partitioned into distinct subpopulations. Naturally, commonly used statistical tests for LD such as Fisher’s exact test or the *χ*^2^ test for association are similarly affected by structure, as they generally assume that HWE holds at one or both loci, i.e., that no structure exists [13,14]. Violating this assumption carries the risk of invalidating *p*-values through inflated Type I error rates and decreased power.

To address these challenges, we introduce a framework to formally test for LD between a pair of loci in a structured population. We work with unphased genotype data from a sample of individuals, conducting a hypothesis test for a quantity called *structural LD* (sLD) – LD that persists after accounting for population structure. To fit arbitrarily complex structure, we utilize an existing model which incorporates covarying allele frequencies among a set of individuals [15–18]. These *individual-specific allele frequencies* (IAFs) are allowed to vary for every SNP–individual pair. This formalization generalizes a wide range of established models, including admixture [16,19,20]. Notably, there is an extensive precedent of IAFs being employed for hypothesis testing, such as in a genome-wide test for association [21], a test for structural HWE (sHWE) [22], and a test for independence among the antecedent populations of the admixture model [18]. IAFs have also been adapted to descriptive statistics, such as a generalized fixation index [16] and more recently an adjusted LD coefficient [23].

Following this paradigm, we develop a hypothesis test that utilizes IAFs for statistical inference while making minimal assumptions about their underlying joint distribution. Because our method assigns *p*-values to locus pairs, we can explicitly quantify the statistical significance of deviations from zero sLD. This enables us to detect correlations *not* driven by structure while remaining robust to spurious, structure-driven LD. The genome-wide distribution of sLD *p*-values also offers a useful criterion for tuning models of population structure, complementing existing tests for sHWE in determining the optimal hyperparameters [22], such as the number of admixed populations, in a model of population structure.

To motivate the sLD test, we briefly summarize here the outcomes of our hypothesis test applied to the Thousand Genomes Project (TGP) [24], which features human genotypes characterized by extensive population structure and recent admixture (further details provided in the Results). In Figure 1, we compare *p*-values from our proposed sLD test for three models of population structure, applying the method to pairs of loci residing on different chromosomes. This is a setting that corresponds to long-range, interchromosomal LD. We implement the sLD test by estimating structure (i.e., IAFs) under an admixture model, where model complexity increases with the number of admixed populations *K*. As *K* increases, the *p*-values from the sLD test progressively converge toward the Uniform(0, 1) distribution, suggesting that the null hypothesis of zero sLD holds when structure is properly modeled. In contrast, applying the sLD test with an unstructured population model (corresponding to *K* = 1 admixed populations) yields significant LD for nearly all locus pairs, demonstrating the pervasive confounding effect of unmodeled structure. This test is equivalent to a traditional test for LD using the sample covariance, which assumes a population to be in HWE. These findings collectively indicate that most interchromosomal LD in this population can be accounted for by a well-fit model of population structure.

**Figure 1.**
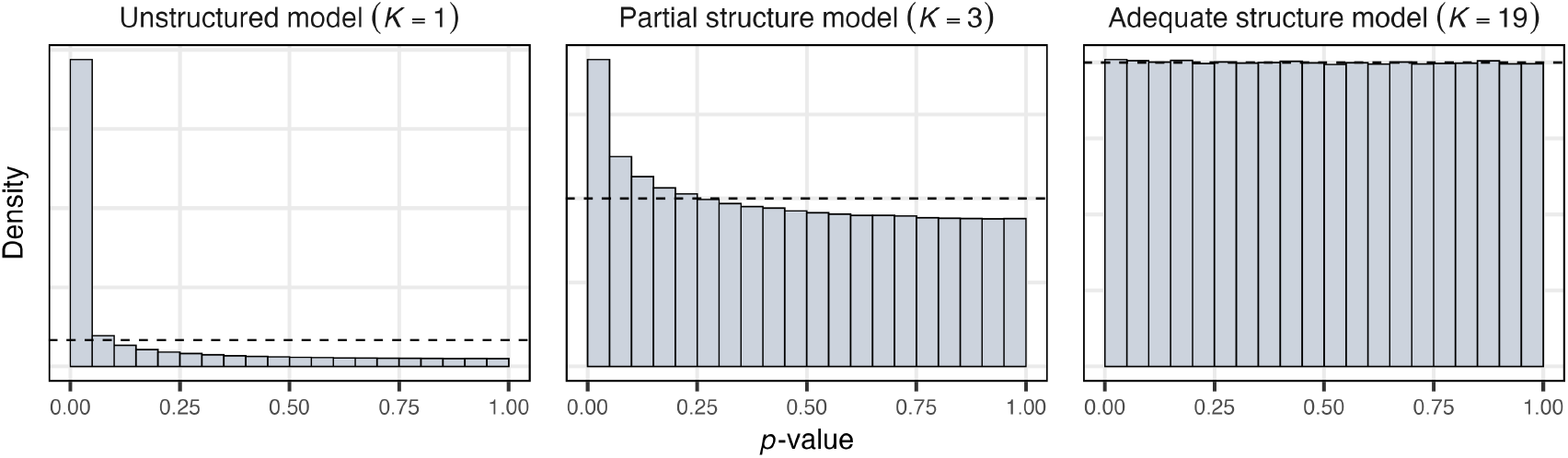
The sLD test applied to the Thousand Genomes Project. Each panel corresponds to a different complexity of the population structure model. Structure was estimated under the admixture model, with *K* denoting the number of admixed populations being modeled. Here, we consider randomly drawn pairs of loci residing on distinct chromosomes. The *p*-values become progressively more uniform as *K* increases and more of the population structure is captured. By contrast, the sLD test employing an unstructured population model (*K* = 1 admixed populations), equivalent to a traditional test for LD, suggests that most loci are in significant LD. The dashed line in each histogram indicates the Uniform(0, 1) density.

We summarize our contributions in this work as follows. In Models and Methods, we provide mathematical details on a model for human genotypes in a population with arbitrarily complex structure, and outline our proposed approach. Results contains simulations demonstrating the utility of the sLD test in the presence of structure. We also show that the framework has the power to detect true instances of structural LD. We then apply the test to human datasets representing highly diverse and structured populations sampled from across the world. We conclude with a Discussion on the applicability of the sLD test to various human genetics settings, caveats, and directions for future development.

## 2 Models and Methods

We begin by summarizing the population model underlying our proposed hypothesis test, called the coancestry model. This model accommodates complex population structure using covarying IAFs. We then introduce our statistical test for sLD. The operating characteristics of the sLD test are investigated in the remainder of this section, with particular emphasis on its desirable theoretical properties when the IAFs are estimated from real data.

### 2.1 A coancestry model of population structure

We have genotype data from *I* individuals at *J* diploid, biallelic loci in a matrix **X**. Let *X*_*ij*_ ∈ {0, 1, 2} denote the measured genotype at locus *i ∈* {1, …, *I*} for individual *j ∈* {1, …, *J*}. For each single nucleotide polymorphism (SNP), a chosen allele is denoted as “1” and the alternative as “0.” Thus, *X*_*ij*_ = 0 and *X*_*ij*_ = 2 represent genotypes homozygous for the “0” and “1” alleles, respectively, and *X*_*ij*_ = 1 equates to a heterozygous genotype. Under the IAF framework, each SNP–individual pair has a potentially unique allele frequency *π*_*ij*_ ∈ (0, 1); these IAFs can be collected into a matrix Π. The simple case of an *unstructured* population corresponds to a given locus having a single allele frequency, with *π*_*i*1_ = *π*_*i*2_ = *⋯* = *π*_*iJ*_. We assume the IAFs to be random parameters, related to measured genotypes via the conditional expectation E[*X*_*ij*_|*π*_*ij*_] = 2*π*_*ij*_.

The coancestry model we employ is an established framework that captures arbitrarily complex population structure, barring close familial relationships, in terms of random, covarying IAFs [17,18]. The term *coancestry* appears in other contexts and is sometimes defined differently; we utilize the model described in [17,18]. This model is defined in terms of moments (i.e., expectations and covariances) for IAFs, which are parameterized with respect to an ancestral population *T*:

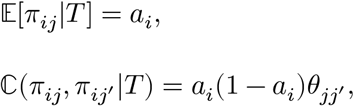

for *i ∈* {1, …, *I*} and *j, j*^′^ ∈ {1, …, *J*}. Here, *a*_*i*_ denotes the frequency of the “1” allele at locus *i* in population *T*, while 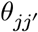 ∈ [0, 1] is the coancestry for individuals *j* and *j*^′^ relative to *T*. For this model, suppose that genotypes have the conditional distribution

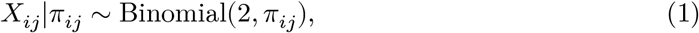

and that the *X*_*ij*_|*π*_*ij*_ are jointly independent, which naturally gives rise to the aforementioned conditional expectation E[*X*_*ij*_|*π*_*ij*_] = 2*π*_*ij*_. Genotypes following the conditional distribution in Equation 1 have been defined to be in *structural Hardy-Weinberg equilibrium* (sHWE) [22]. This distribution results in the following covariances for genotypes:

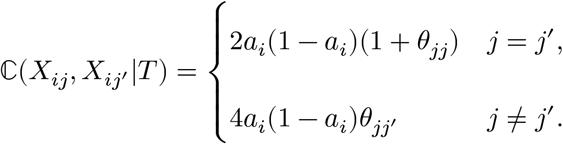

These covariances are widely used in modeling relatedness and structure within populations when working with genotype data [17,18].

For our inference framework, we view the IAF matrix Π as a specific realization of this coancestry model. In a given study, we sample individuals from a study population *S*, which itself is derived from an ancestral population *T* according to an arbitrary evolutionary process satisfying the moments of the coancestry model (Figure 2). We express this as Π_*S*_ *∼* ℒ_*π*_, where Π_*S*_ denotes the set of *all* IAFs within *S*, and ℒ_*π*_ is the evolutionary process leading from *T* to *S*. We do not specify an explicit form for ℒ_*π*_, only requiring that its moments are consistent with the coancestry model. It follows that 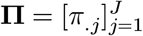 is essentially a random sample from Π_*S*_, where *π*_.*j*_ ∈ ℝ^*I*^ denotes the column vector of individual *j*’s IAFs for all *I* SNPs. Here, we let [*⋅*] denote concatenation across columns, such that Π is composed of a set of *J* total columns, each representing a given individual’s IAFs across *I* loci.

**Figure 2.**
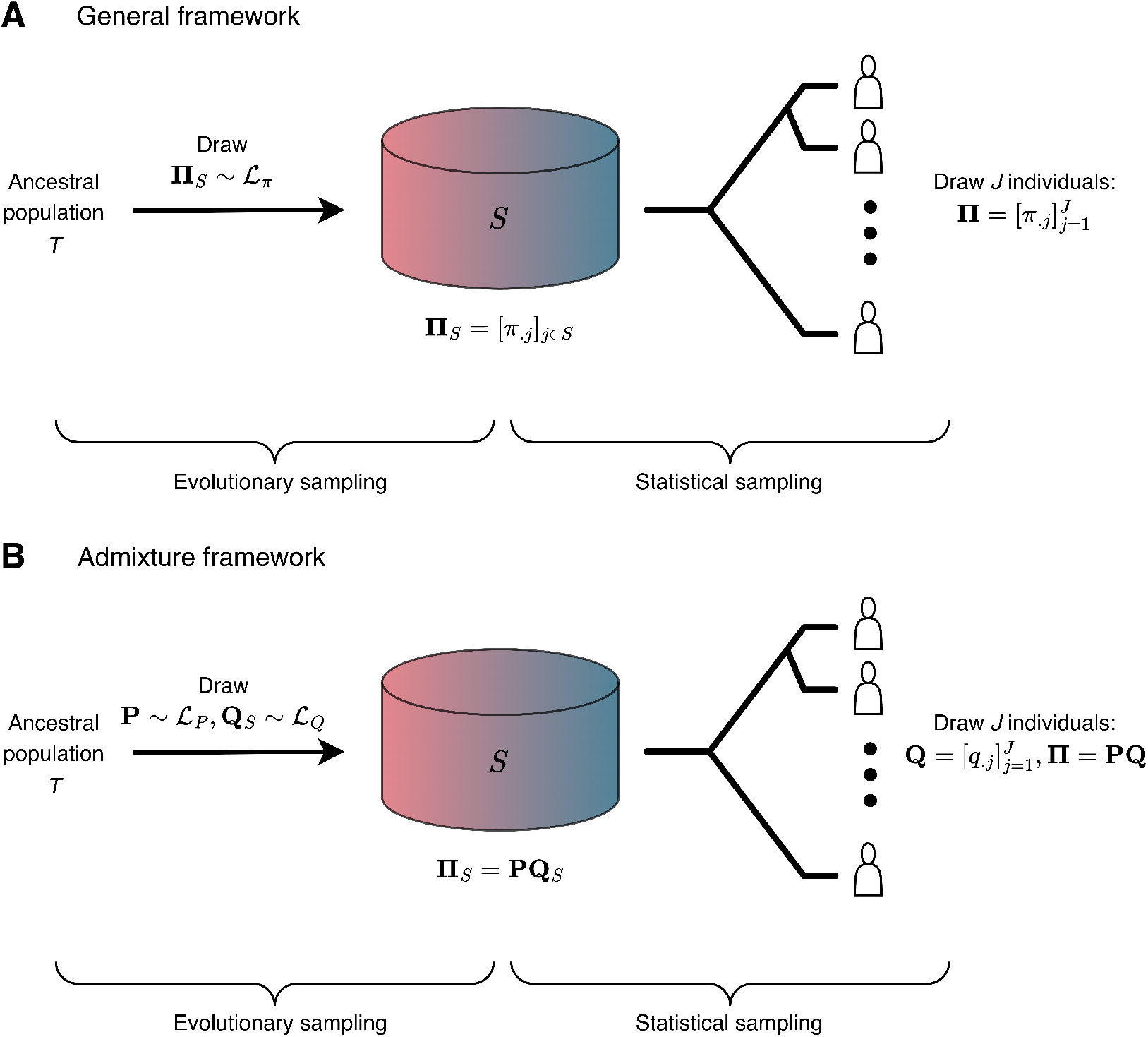
The joint distribution underlying IAFs is characterized by two layers of random sampling: evolutionary sampling leading from an ancestral population *T* to the study population *S*, and statistical sampling of *J* individuals from *S*. (**A**) The general coancestry framework supposes that the study population *S* is itself a random draw from some evolutionary process originating from *T*. We then randomly sample *J* individuals from *S* to yield a matrix of IAFs Π. (**B**) The admixture model can be viewed as a specific example of this framework. In particular, all individuals within *S* derive their genetic ancestries from *K* admixed populations, with varying proportions according to their respective admixture vectors. The allele frequencies of the admixed populations are contained within a matrix **P**, while the admixture vectors are collected to form the columns of a matrix **Q**; these *q*_1_, *q*_2_, …, *q*_*J*_ are a representative sample of a larger, population-wide set of IAFs **Q**_*S*_. The matrix of IAFs therefore has the form Π = **PQ**.

A well-developed model that outlines an explicit generative process for IAFs is the admixture model, which posits that each individual derives some proportion of their genetic ancestry from each of *K* non-overlapping admixed populations. The admixture model is a useful example of the coancestry model for two reasons. First, it formally instantiates the general framework outlined previously and specifies an explicit sampling mechanism for IAFs (Figure 2). Secondly, it offers a tractable approach for estimating Π from real genotypes. Under this model, IAFs are a weighted average of admixed population allele frequencies

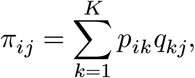

where 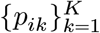 are the allele frequencies at locus *i* in each admixed population and 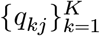 are the admixture proportions for individual *j*, denoting the share of their genetic ancestry from each population. In matrix notation, this corresponds to the factorization Π = **PQ**, with **P** and **Q** denoting *I* × *K* and *K* × *J* matrices of admixed population allele frequencies and admixture proportions, respectively. Here, the number of admixed populations *K* is effectively a hyperparameter that tunes the complexity of structure modeled.

As a specific instance of the coancestry model, the admixture model can also be framed with respect to a study population *S* and an ancestral population *T*. The study population *S* is characterized by a set of admixed population allele frequencies **P** randomly drawn from a distribution ℒ_*P*_, and a population-wide collection of admixture proportions **Q**_*S*_ from a distribution ℒ_*Q*_. Both ℒ_*P*_ and ℒ_*Q*_ represent random processes leading from *T* to *S*. The admixed population allele frequencies **P** are a component shared across all individuals within *S*, including those in our study. Conversely, the admixture proportions in **Q**_*S*_ are individual-specific, with each admixture vector *q*_.*j*_ capturing the unique ancestral composition of individual *j*. Within a given study, the admixture proportions for *J* individuals 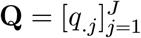 constitutes a representative sample of **Q**_*S*_. Technical details regarding the admixture model and the estimation of IAFs from real data are contained within Appendix B.

### 2.2 A test for structural linkage disequilibrium

To demonstrate how population structure induces LD, let us observe the effects of structure on the covariance between genotypes at two different loci. Using the law of total covariance, we can decompose the marginal covariance ℂ_*S*_(*X*_*a*_, *X*_*b*_) between a pair of genotypes *X*_*a*_ and *X*_*b*_ for an arbitrary individual in *S* as follows:

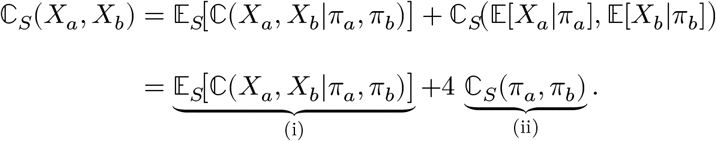

As such, this marginal covariance is driven by two distinct components, labeled (i) and (ii). Note that while we use the coancestry model, which conditions moments on an ancestral population *T*, we omit this notation for brevity. The component in the first term, ℂ (*X*_*a*_, *X*_*b*_|*π*_*a*_, *π*_*b*_), is the conditional covariance between genotypes *X*_*a*_ and *X*_*b*_ given their respective IAFs *π*_*a*_ and *π*_*b*_. We call ℂ (*X*_*a*_, *X*_*b*_|*π*_*a*_, *π*_*b*_) the *structural LD* (sLD) parameter to emphasize that it is the residual covariance that persists after explicitly accounting for population structure via IAFs. The second term, ℂ_*S*_(*π*_*a*_, *π*_*b*_), captures the covariance between allele frequencies at loci *a* and *b* within *S* (given *T*), a quantity that is generally nonzero for a structured population. Therefore, even if the sLD parameterℂ (*X*_*a*_, *X*_*b*_|*π*_*a*_, *π*_*b*_) = 0, as is the case for loci that are conditionally independent given their IAFs, the marginal covariance on the left-hand side, ℂ_*S*_(*X*_*a*_, *X*_*b*_), can still be nonzero if the IAFs have a nonzero covariance, i.e., ℂ_*S*_(*π*_*a*_, *π*_*b*_) ≠ 0. It follows that a hypothesis test utilizing the marginal covariance ℂ_*S*_(*X*_*a*_, *X*_*b*_) may identify structure-driven LD when structure exists in the study population *S*. We focus on the sLD parameter to assess whether a pair of loci are independent after explicitly conditioning on their covarying allele frequencies.

Our method aims to perform a hypothesis test of the sLD parameter ℂ (*X*_*a*_, *X*_*b*_|*π*_*a*_, *π*_*b*_), evaluating whether it is zero or nonzero. To enable the forthcoming discussion, we will first extend the notion of IAFs to *individual-specific haplotype frequencies* (IHFs). We use the term *haplotype* in a functional sense to denote a pair of alleles at distinct loci originating from a single parental source (maternal or paternal), without requiring physical contiguity along a given chromosome. As such, let *H*_*h,j*_ ∈ {0, 1, 2} denote the count of haplotype *h ∈* {*ab, aB, Ab, AB*} in individual *j* at two distinct loci *a* and *b*, and *η*_*h,j*_ its corresponding frequency. Here, lowercase and uppercase letters represent the “1” and “0” alleles, respectively, in accordance with the notation for the coancestry model. We model haplotype counts as being drawn according to

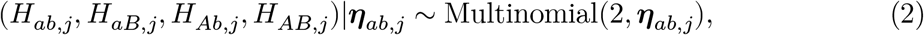

for all *j ∈* {1, …, *J*}, where *η*_*ab,j*_ := (*η*_*ab,j*_, *η*_*aB,j*_, *η*_*Ab,j*_, *η*_*AB,j*_) denotes a vector of IHFs constrained according to *π*_*aj*_ = *η*_*ab,j*_ + *η*_*aB,j*_ and *π*_*bj*_ = *η*_*ab,j*_ + *η*_*Ab,j*_. Note that the four IHF components must also sum to one. Individual *j*’s genotypes at loci *a* and *b* can be written as sums of haplotype counts:

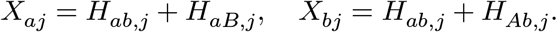

In the trivial case where *a* = *b*, the formulation above reduces to the Binomial distribution in Equation 1, which treats the alleles at a single locus as independently and identically distributed (i.i.d.) Bernoulli random variables conditional on the IAFs. As such, we generalize this idea by treating genotype *pairs* as sums of i.i.d. maternal and paternal haplotypes. It should be mentioned that the genotypes comprising any such pair still marginally follow the Binomial distribution (see Appendix A for a proof).

The haplotype frequency *η*_*ab,j*_ serves as the parameter of interest in our hypothesis test.

The test identifies its signature through a product of mean-centered genotypes for each individual as

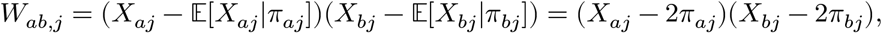

where we utilize the fact that E[*X*_*ij*_|*π*_*ij*_] = 2*π*_*ij*_ for any *i, j* in the coancestry model we employ. The expectation of this quantity equals the conditional covariance for individual *j*, so that

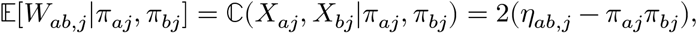

where the second equality follows from the Multinomial distribution in Equation 2. We formally test the hypotheses:

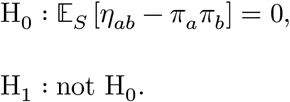

Note that there are no indices on the genotypes and IAFs corresponding to any particular individual *j*. This is because our test concerns the average covariance structure in *S* rather than the covariance of a given individual; by treating the *J* individuals in our study as a random sample of *S*, we can test whether the individual-specific covariance *η*_*ab*_ − *π*_*a*_*π*_*b*_ is nonzero on average for individuals randomly sampled from *S*. Here, the expectation E_*S*_ emphasizes that we are conducting inference on this particular study population *S* – a specific realization of some random evolutionary process.

The sLD test is implemented as follows. 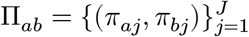 denote the set of true IAFs at loci *a* and *b* for all *J* individuals. When working with real data, we do not observe the true IAFs Π_*ab*_, so we instead form an estimate 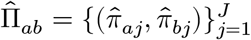. The *theoretical* sLD test statistic, which employs the true IAFs, is defined as

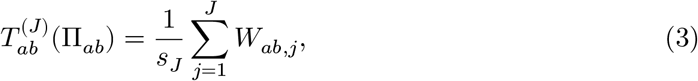

where the denominator accounts for the variance under the null hypothesis:

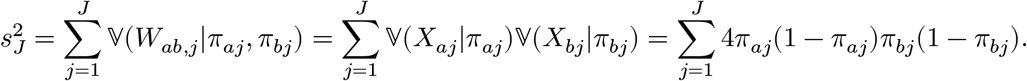

Note that the terms on the right-hand side of Equation 3 are also conditioned on Π_*ab*_, though we omit this for brevity. The variance term 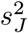 leverages the fact that V(*X*_*ij*_|*π*_*ij*_) = 2*π*_*ij*_(1 − *π*_*ij*_) when Equation 1 holds, and also a statistical property proved in Appendix A. In practice, we implement the test using the estimated 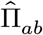, replacing the true IAFs *π*_*aj*_, *π*_*bj*_ with their corresponding estimates 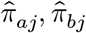 wherever they appear in the test statistic. As such, we call 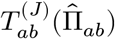 the *implemented* test statistic. The asymptotic null distributions of the theoretical and implemented test statistics are detailed in Theoretical operating characteristics of the sLD test.

The sLD test is designed to be applied genome-wide across many pairs of loci, without restrictions on pairs within or across chromosomes. Its primary objective is to identify pairs of loci with significant structural LD. Importantly, its implementation requires only observed genotypes, from which estimated IAFs 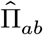are obtained. A wide range of methods exists for estimating IAFs; a well-known and flexible parameterization is the admixture model. While we use a semiparametric estimator of the admixture model [20] in our analyses, the sLD test accommodates alternative admixture estimators and extends to a broader coancestry framework that contains admixture as a special case. Furthermore, although the hypotheses being tested are framed in terms of the conditional covariance between loci *a* and *b*, we can indeed view the sLD test as a test of conditional *independence* between loci *a* and *b*. We detail this result in Appendix A.

### 2.3 Theoretical operating characteristics of the sLD test

We examine the operating characteristics of the sLD test in the regime of a large number of individuals *J*, deriving the aysymptotic null distribution of the theoretical sLD test statistic 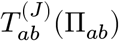 (i.e., when the IAFs are known). We state a simple condition under which 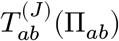 converges in distribution to a standard Normal random variable in Theorem 2.1 below.

#### Theorem 2.1.

*Let π*_*aj*_, *π*_*bj*_ ∈ [*ϵ*, 1 −*ϵ*] *for some arbitrarily small ϵ ∈* (0, 1/2] *for all j. Under the null hypothesis and the coancestry model, the theoretical sLD test statistic in Equation 3 has convergence*

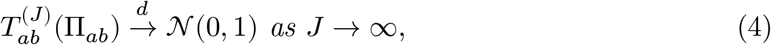

*where* 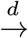 *denotes convergence in distribution*.

The proof of Theorem 2.1 is provided in Appendix A, and follows from a specific version of the Central Limit Theorem [25]. Intuitively, when the IAFs do not exactly equal 0 or 1 and the sample size is large, the theoretical sLD test statistic has a well-defined asymptotic distribution under the null.

While Theorem 2.1 applies to the theoretical test statistic 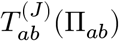, we do not observe the true IAFs Π_*ab*_ in practice. Our goal is to extend this convergence result to be informative about the implemented test statistic 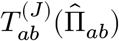, which utilizes estimated IAFs 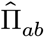. Here, we provide theoretical conditions ensuring that 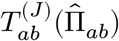 itself converges to the standard Normal distribution.

#### Corollary 2.1.

*Consider the implemented test statistic* 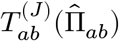, *which takes the form in Equation 3 but replaces the set of true IAFs* Π_*ab*_ *with their estimates* 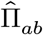. *If*

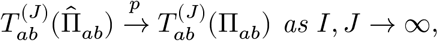

*where* 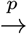 *denotes convergence in probability, then*

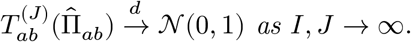

The result above follows directly from an application of Slutsky’s theorem [26]. The requirement that *I* → ∞ (jointly with tbe number of individuals *J* → ∞) highlights how our chosen framework leverages a large number of SNPs *I* across the genome to achieve consistency [27], estimating the entire matrix of IAFs Π simultaneously. This result is appealing because it establishes that the implemented test statistic 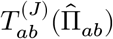, which we evaluate in practice, admits an analytical asymptotic null distribution under certain conditions. By contrast, a related hypothesis test for sHWE [22] utilizes an empirical null distribution, repeatedly sampling from a fitted model of population structure to account for additional variability induced by the process of estimation. Our analyses in the Results support the notion of 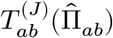 following the Normal(0, 1) distribution under the null hypothesis, though the aforementioned empirical null [22] offers a useful alternative if the conditions in Corollary 2.1 are not satisfied.

## 3 Results

We first apply the sLD test to simulated genotypes to validate its theoretical properties. We then analyze three publicly available studies – the Thousand Genomes Project (TGP) [24], the Human Genome Diversity Project (HGDP) [28], and the Human Origins (HO) study [29]. These resources feature diverse genotype data from across the world containing substantial population structure, providing challenging and informative settings in which to evaluate the sLD framework.

### 3.1 Simulation studies

We first use simulated data to show that population structure induces spurious, structure-driven LD, and demonstrate that our method accurately accounts for such. All pairs of loci in these simulations follow the null hypothesis of no structural LD. However, we may observe spurious LD for any given pair of loci due to structure; not adjusting for the presence of structure can lead to challenges in disentangling structure-driven LD versus LD generated via other sources. We simulate genotypes under an admixture model wherein admixture proportions reflect a random walk along a one-dimensional geography [16]. This model emulates spatial population structure and represents a particularly challenging instance of structure to estimate and analyze [16,20]. In Figure 3, we show that the sLD test yields null *p*-values closely following the Uniform(0, 1) distribution when structure is accurately modeled. This contrasts with the sLD test using an unstructured population model (an admixture model that reduces to *K* = 1 admixed populations), wherein all IAFs at a given locus are set equal to a common allele frequency for all individuals. The formulation here is equivalent to a test for LD using the sample covariance, so we will refer to the *K* = 1 sLD test as the traditional test for LD hereafter. In this *K* = 1 case, the *p*-values are strongly skewed towards zero, meaning we would identify nearly all locus pairs as exhibiting significant LD. We provide technical details regarding these simulations in Appendix B.

**Figure 3.**
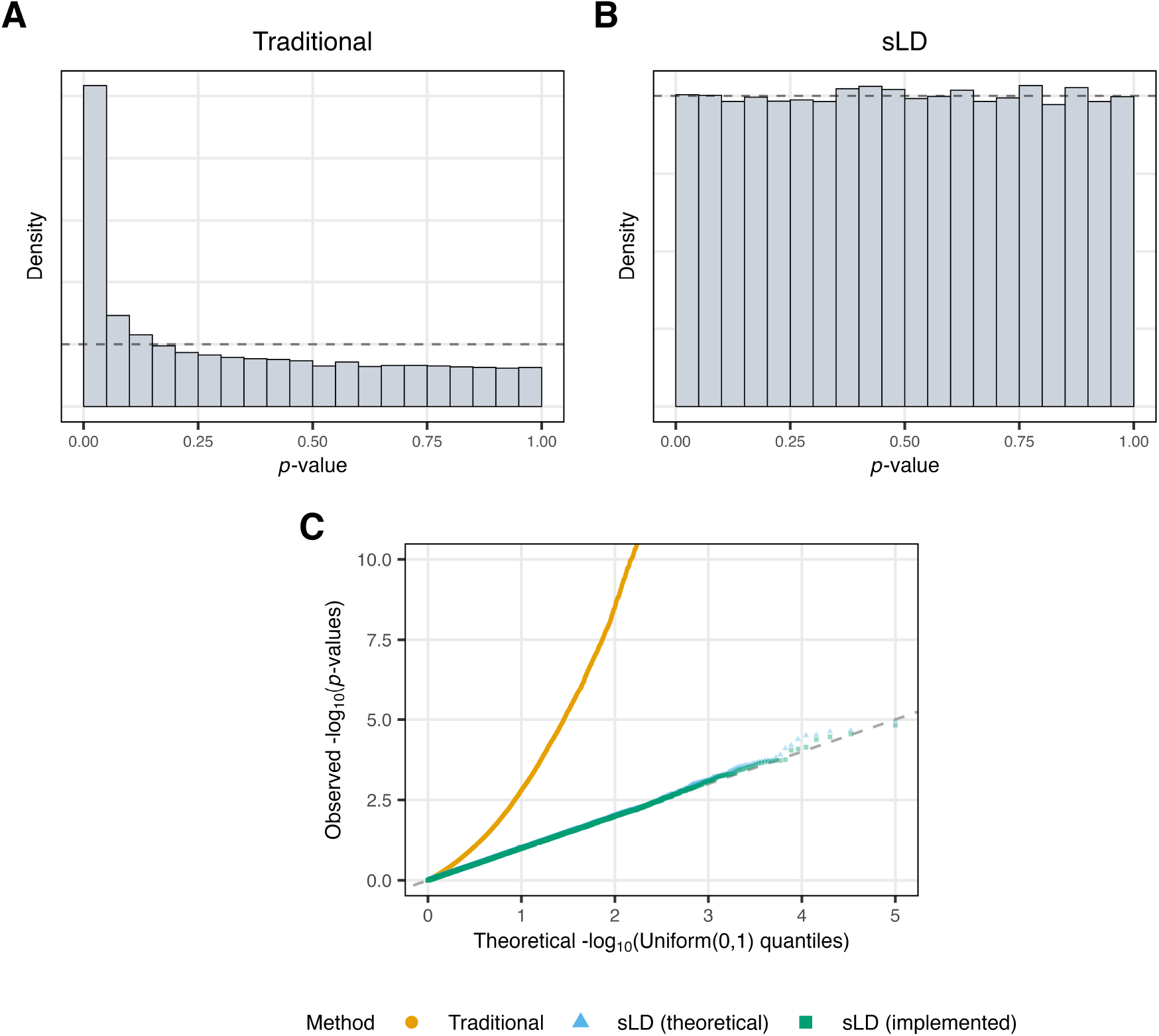
Simulation results for locus pairs drawn from an admixture model under the null hypothesis of no structural LD. (**A**) Histogram of *p*-values corresponding to the traditional test for LD (*K* = 1 sLD test). (**B**) Histogram of *p*-values for the sLD test with an accurate model of population structure, employing the theoretical test statistic 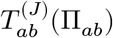 with known IAFs. The dashed line in both histograms indicates the Uniform(0, 1) density. (**C**) Quantile-quantile plot of null *p*-values for the traditional test for LD (orange), the sLD test with the true IAFs (blue), and the sLD test utilizing estimated IAFs (green).

Our simulations also support the theoretical results in Theoretical operating characteristics of the sLD test regarding the asymptotic distribution of the implemented test statistic 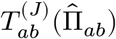. Specifically, for representative study sizes (e.g., *I ∼* 10^5^ − 10^6^ loci and *J ∼* 10^2^ individuals), the sLD test yields null *p*-values that follow the Uniform(0, 1) distribution even when the IAFs are estimated (Figure 3). We also evaluate the theoretical Normal(0, 1) null distribution by comparing it with an empirical null distribution obtained using the parametric bootstrap approach of ref. [22] (see Appendix B). We perform this comparison using genotypes simulated from IAFs estimated on real data to determine whether the Normal(0, 1) null is appropriate when analyzing these datasets. We find that the Normal(0, 1) is a reasonable approximation to the distribution of null test statistics for all three human studies.

We conclude our simulation studies by demonstrating that the sLD test has statistical power to detect structural LD. Using a forward-generating probabilistic framework outlined in Appendix A, we simulate a combination of genotype pairs in structural LD versus not in structural LD. We evaluate statistical power at different levels of structural LD by adjusting the value of an individual-specific LD parameter *δ*_*ab,j*_. To account for differing allele frequencies across SNP–individual pairs, we parameterize *δ*_*ab,j*_ as a function of the IAFs *π*_*aj*_, *π*_*bj*_ and a correlation parameter *ρ*_*ab*_ shared across all individuals in a given study. Intuitively, larger values of *ρ*_*ab*_ correspond to stronger structural LD between loci *a* and *b* within the study population. We provide details on this parameterization in Appendix B. Figure 4 summarizes these simulations, showing that the sLD test implemented with a structured population model has comparable power to the traditional test for LD (the *K* = 1 sLD test). By contrast, control of the False Discovery Rate (FDR) [30] at the desired threshold is achieved only when structure is modeled. This result holds for both the theoretical test statistic 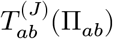 and the implemented test statistic 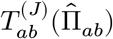, employing the true IAFs and estimated IAFs, respectively. In other words, the sLD test, when structure is accurately modeled or estimated, is able to identify structural LD as well as the traditional test for LD, but remains robust to spurious LD. As with previous simulations, implementation details are documented in Appendix B.

**Figure 4.**
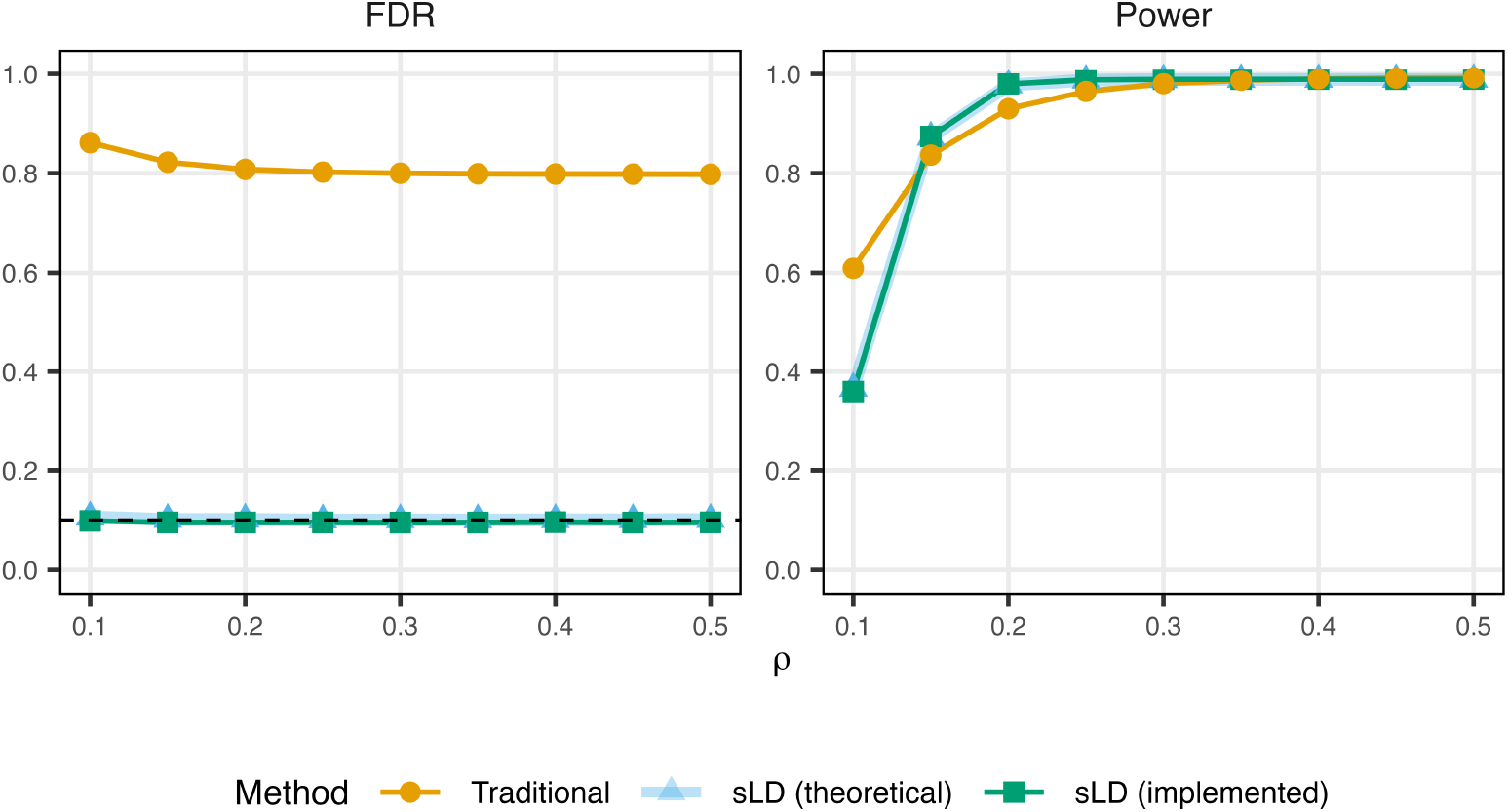
Simulations evaluating the statistical power of tests for LD, comparing the traditional test for LD (orange), the sLD test with the true IAFs (blue), and the sLD test employing estimated IAFs (green). We simulate genotypes under a mixture of null (no structural LD) and non-null (structural LD) scenarios. While each test type has comparable statistical power, control of the FDR is maintained only under the sLD test when population structure is modeled or estimated. The dashed line indicates a nominal FDR threshold of 0.1.

### 3.2 Analyses of human studies

We apply the sLD test to the TGP, HGDP, and the HO studies. Details regarding the processing of these datasets and the estimation of IAFs are contained within Appendix B. In brief, we use a semiparametric estimator of the admixture model, with *K* denoting the number of admixed populations modeled. We first consider pairs of loci on distinct chromosomes to ascertain whether interchromosomal (long-range) LD can be accounted for by population structure. Figure 5 demonstrates that sLD test *p*-values for pairs spanning distinct chromosomes approximately follow the Uniform(0, 1) distribution for all three datasets when population structure is accurately modeled and estimated. For each dataset, we randomly draw 2 million interchromosomal locus pairs for computational feasibility. We apply the entropy-based procedure for selecting *K* developed for the related sHWE test [22], which evaluates a range of *K* values by measuring how close their corresponding *p*-value distributions are to the Uniform(0, 1) distribution. We apply this criterion to both sHWE and sLD test *p*-values, reporting their respective choices of *K* (Figure 5). We note that while the original algorithm for the sHWE test omitted the lowest *p*-value interval near zero when calculating entropy due to a large, persistent peak here, the interchromosomal sLD *p*-values show a smaller peak near zero. Because of this, we chose to include the entire range [0, 1] when calculating entropy for sLD *p*-values. Collectively, our results indicate that most long-range LD can be accounted for by well-fit models of structure; by contrast, assuming an unstructured population model leads to a large number of *p*-values near zero for all three studies.

**Figure 5.**
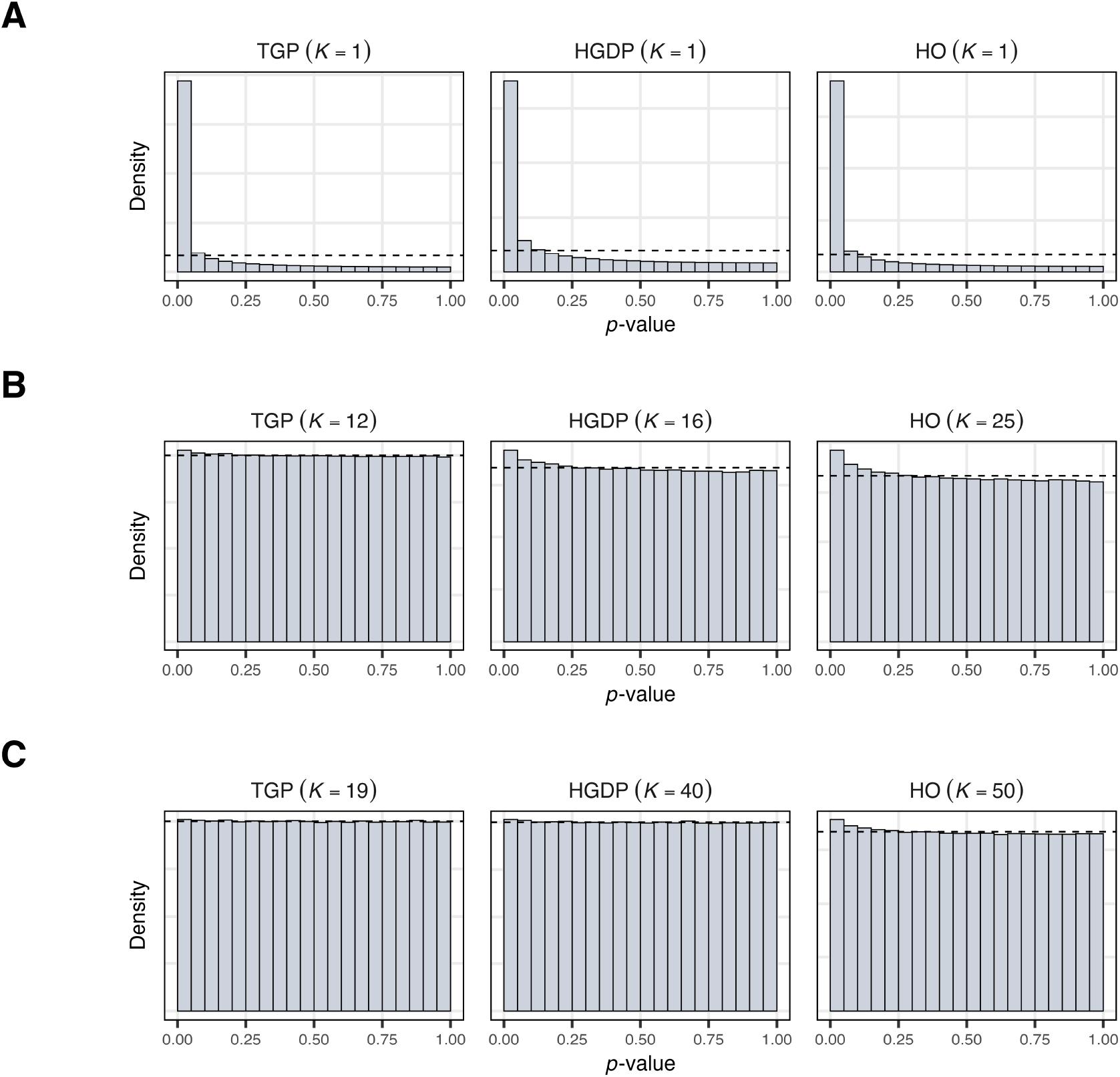
Histograms of sLD *p*-values for locus pairs spanning different chromosomes for the TGP, HGDP, and HO studies. The dashed line in each histogram indicates the Uniform(0, 1) density. (**A**) Histograms corresponding to the traditional test for LD (*K* = 1 sLD test) for each study. (**B**) Histograms with *K* chosen by the entropy criterion applied to sHWE test *p*-values. (**C**) Histograms with *K* chosen by the entropy criterion applied to sLD *p*-values directly.

We observe that the number of admixed populations *K* required for the sLD *p*-values to become uniform is larger than the value selected by the sHWE test for each dataset. We expect this to be the case, as these *p*-values are driven by the joint distributions of pairs of loci, which may be more complex than the marginal distributions of individual loci. We also note that a larger value of *K* is necessary for the HGDP and HO datasets relative to HGDP to display uniform *p*-values. This may be attributed to instances of more recent admixture within these populations [29] as well as finer-grained structure, so that larger numbers of admixed populations are necessary to capture this increased complexity. This is especially evident for the HO study, as has been previously observed [22]; indeed, a small peak near zero is maintained even for a larger number of admixed populations *K* than the other two studies. As with the simulation studies, Appendix B contains implementation details and our sampling processes for locus pairs under different scenarios, such as interchromosomal versus intrachromosomal LD.

We then apply the sLD test to locus pairs within a given chromosome. Comparing the *p*-values under the sLD test (with a structured population model) against the traditional test for LD, we find that *p*-values at close physical distances are very small for both test types.

By contrast, *p*-values increase more sharply with growing physical distances when structure is modeled (Figure 6). The structure model we implement for each dataset corresponds to an admixture model with *K* chosen via the sHWE test [22]. These results suggest that the statistical significance of LD decays at a faster rate with physical distance when structure is adequately captured; LD also has a less significant baseline at very long ranges (*∼* 10^5^ − 10^6^ base pairs). We visualize this same pattern when focusing on specific regions in the genome. We investigated the HLA class II region on the short arm of chromosome 6, which contains alternating tracts of suppressed and elevated recombination [31]. Looking at the TGP, we see that the sLD test identifies short-range blocks as exhibiting statistically significant LD, much like the traditional test for LD, while longer ranger LD and LD between blocks is less significant after accounting for population structure (Figure 7).

**Figure 6.**
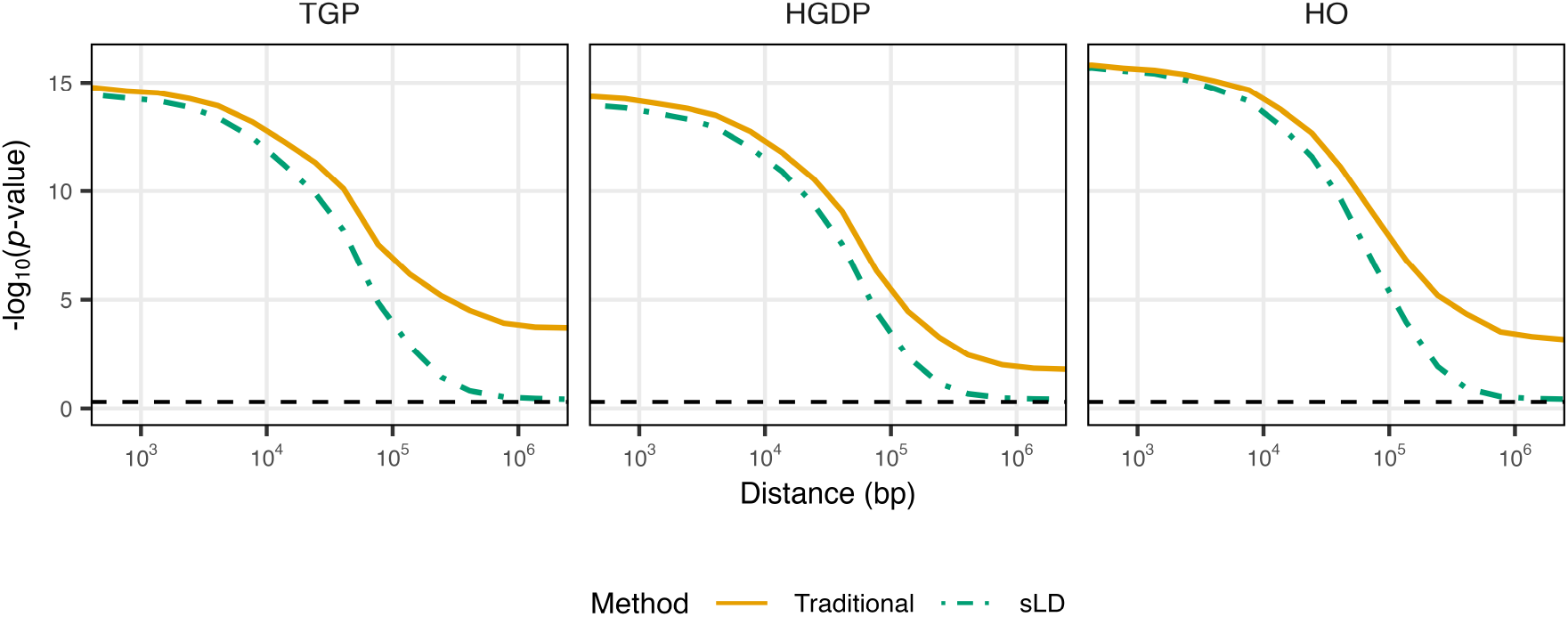
Plots of mean – log_10_ (*p*-values) against physical distance between loci for the TGP, HGDP, and HO studies. For each dataset, we sample pairs of loci residing on a given chromosome, and compute *p*-values corresponding to both the traditional test for LD (orange) as well as the sLD test with a structured population model (green). Locus pairs were sampled from all 22 autosomes. We truncate − log_10_ (*p*-values) at 16 to accommodate *p*-values that either equal or approach zero. The dashed line marks the position of − log_10_ (0.5), which represents the median *p*-value under a Uniform(0, 1) distribution.

**Figure 7.**
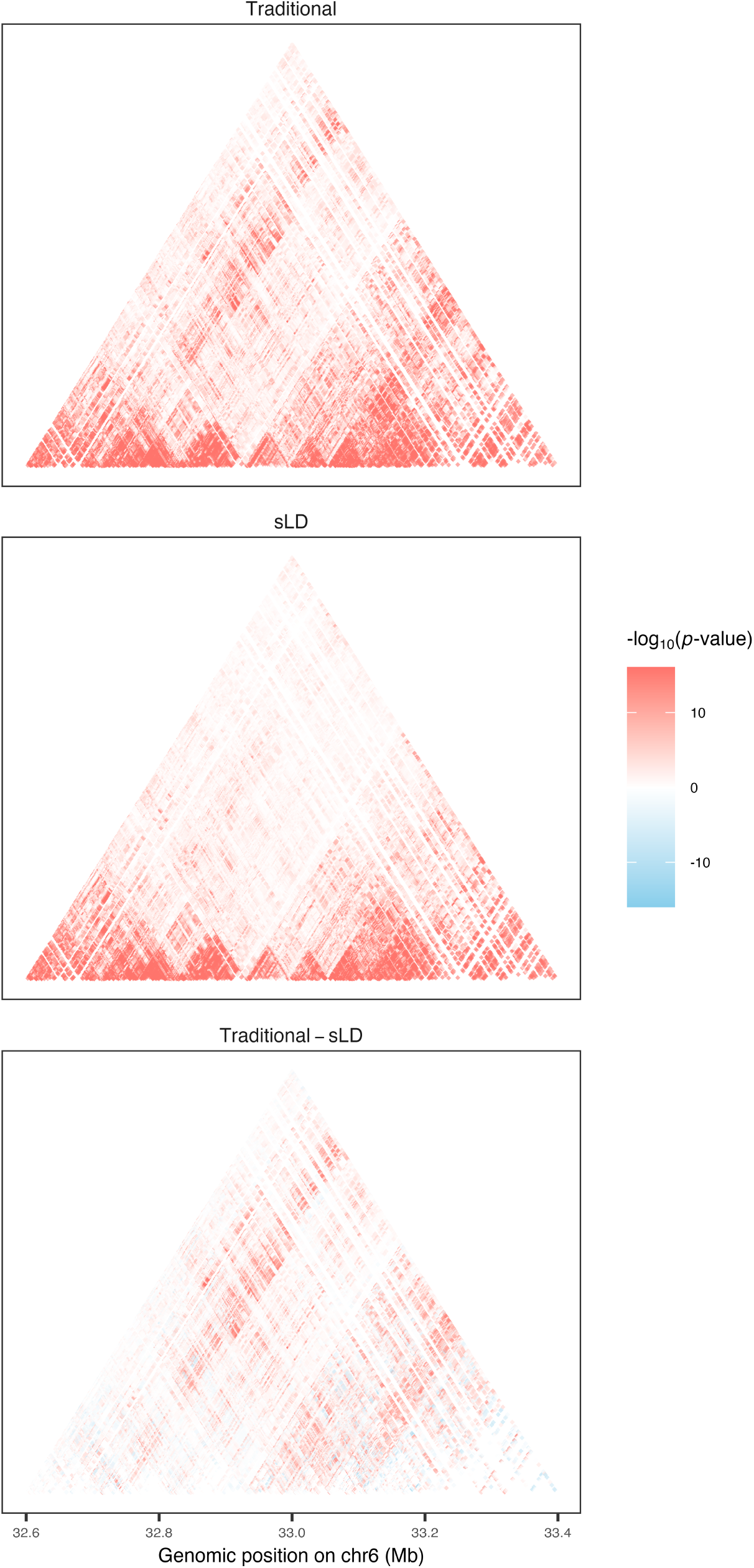
Heatmaps of sLD *p*-values for locus pairs within the HLA Class II region on chromosome 6, computed from the TGP. Here, we plot − log_10_ (*p*-values) for the sLD test and the traditional test for LD. We also plot the difference in − log_10_ (*p*-values) between the two test types.

## 4 Discussion

LD plays an essential role in genomics research, underpinning analyses such as association studies, and significant efforts have been made to characterize it on a genome-wide basis. Population structure is pervasive in human populations and induces linkage disequilibrium even when no other factors driving LD are present; LD arising from structure is sometimes referred to as spurious LD. In light of this, we developed the sLD test, a method that tests for LD between a pair of loci in a structured population. Specifically, we test for a quantity termed structural LD, which measures the conditional covariance between locus pairs after accounting for population structure and relatedness, ensuring that any remaining LD is not due to structure.

We applied the sLD test to three human studies: the Thousand Genomes Project, Human Genome Diversity Project, and the Human Origins study. Our results suggest that interchromosomal LD can be accounted for by well-fit models of population structure, in that sLD test *p*-values for pairs of loci spanning different chromosomes are approximately Uniform(0, 1) distributed after modeling structure. By contrast, testing for LD *without* accounting for structure would indicate that nearly all interchromosomal locus pairs are in statistically significant LD. We also show that within individual chromosomes, the sLD framework mitigates the statistical significance of LD driven by structure across long distances, while LD at small base pair distances remains highly significant, consistent with genetic linkage at such close proximity.

Our method builds upon an existing population model for genotypes called the coancestry model [17,18]. As such, it inherits the flexibility of the coancestry model, supporting a wide range of parameterizations of population structure. We implement an estimator of the admixture model for our analyses, though we emphasize that the sLD test is compatible with a more general framework that includes admixture as a specific case. Importantly, we can validate the model fit of a given structure model on a locus-by-locus basis, using existing methods like the related sHWE test [22]. The sLD test also does not require phased genotypes, and does not explicitly involve the estimation of haplotypes or haplotype frequencies.

Because the sLD test statistic has a theoretical Normal(0, 1) distribution under the null hypothesis, the method is computationally efficient to implement. This property obviates the need to estimate an empirical null distribution, which can require extensive permutation or resampling. However, the Normal(0, 1) null is reliant on certain assumptions, which should be validated for a given study and population model. More generally, it is important to confirm that the specific model of population structure and relatedness is reasonable for the study. If the assumptions for the theoretical Normal(0, 1) null are not satisfied, the procedure for simulating an empirical null distribution described in [22] offers an effective alternative.

There are several additional considerations when implementing the sLD test. The sLD test requires identifying a specific set of locus pairs to investigate, as it is generally not feasible to apply the test to every possible pair of loci for a given study. For example, if we had *I* = 10^6^ loci, testing all unique pairs corresponds to 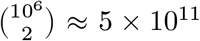 hypothesis tests. Investigators must additionally choose a specific significance threshold for locus pairs, such as controlling the FDR, the expected number of false positives, or the local FDR; the appropriate choice may depend on the goals of the study. For any significant structural LD detected by the sLD test, it also remains open to pinpoint the specific mechanism underlying it outside of structure. Some possibilities include genetic linkage and selective pressures favoring a specific set of alleles at different loci (epistatic selection).

While we implemented the sLD test using admixture, future work could utilize other implementations of the coancestry model, such as a model of structure built upon continuous variation along a spatial axis. Another important avenue of investigation could be in the design of new genotyping arrays for diverse human cohorts, and in particular the total human population; these are settings where population structure is likely to be widespread. Because some signals of LD may represent spurious LD due to structure, using structural LD could help optimize the selection of marker SNPs and ensure uniform coverage of the genome. The sLD test may also prove valuable for the investigation of certain population-level processes. For example, a pair of interchromosomal loci in strong structural LD may represent a signal of epistatic selection. Adapting the sLD framework into a specialized scan for natural selection represents an interesting direction for future research.

## Data Availability

A reproducible analysis document containing additional technical details, as well as implementations of simulations and real data analyses, can be found at https://github.com/StoreyLab/sLD-manuscript-analysis. An R package implementing the methods can be found at https://github.com/StoreyLab/sLDtest.

## Study Funding

This work was supported in part by United States National Institutes of Health (HG006448 to JDS). The funders had no role in study design, data collection and analysis, decision to publish, or preparation of the manuscript.

## Conflict of Interest

The authors declare no conflicts of interest.

## A Supplementary Theory and Methods

### A.1 A forward-generating probability model of genotypes in structural LD

We outline a forward-generating probability model for pairs of genotypes in structural LD. We use this model to conduct the statistical power simulations in the Results. This model builds on the individual-specific haplotype frequency (IHF) formulation in Equation 2 (see A test for structural linkage disequilibrium), restated here for convenience:

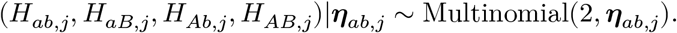

The *H*_*h,j*_’s are haplotype counts as defined in Equation 2, with *h ∈* {*ab, aB, Ab, AB*} denoting one of the four possible haplotypes; the vector *η*_*ab,j*_ := (*η*_*ab,j*_, *η*_*aB,j*_, *η*_*Ab,j*_, *η*_*AB,j*_) contains their corresponding IHFs. The IHFs are constrained to sum to one and relate to individual-specific allele frequencies (IAFs) via *π*_*aj*_ = *η*_*ab,j*_ + *η*_*aB,j*_ and *π*_*bj*_ = *η*_*ab,j*_ + *η*_*Ab,j*_. We can then form genotypes after drawing haplotype counts, with *X*_*aj*_ = *H*_*ab,j*_ + *H*_*aB,j*_ and *X*_*bj*_ = *H*_*ab,j*_ + *H*_*Ab,j*_.

To produce genotypes in structural LD, we can parameterize IHFs as follows:

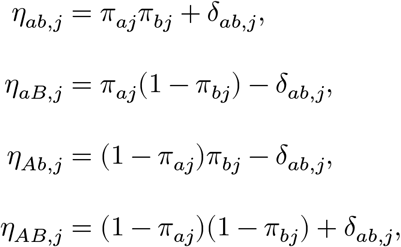

where *δ*_*ab,j*_ denotes an individual-specific LD parameter for loci *a* and *b* in individual *j*. Note that these IHFs satisfy the constraints outlined previously, namely that they sum up to one, and we have that

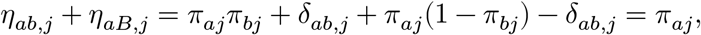

with an analogous result holding for *π*_*bj*_. Here, *δ*_*ab,j*_ plays a similar role to the classical LD coefficient *D*_*ab*_ for a sample of individuals [3], only that it is specific to a particular individual. The absence of structural LD between loci *a* and *b* equates to the scenario where E_*S*_[*δ*_*ab*_] = 0; by contrast, structural LD exists when E_*S*_[*δ*_*ab*_] ≠ 0. We also require *δ*_*ab,j*_ to be feasible for the pair of allele frequencies *π*_*aj*_, *π*_*bj*_ so that it is bounded by the minimum and maximum possible values of the LD coefficient for that pair, with

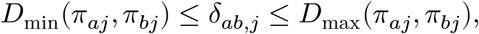

where

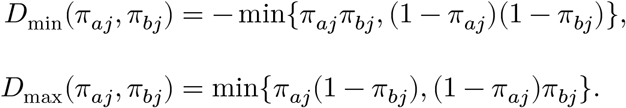

Bounding the individual-specific LD term *δ*_*ab,j*_ in this manner ensures that each of the IHFs will be valid haplotype frequencies confined to the interval [0, 1].

Under this forward-generating probability model, we can write each of the IHFs *η*_*ab,j*_ at loci *a* and *b* in terms of a pair of individual-specific allele frequencies *π*_*aj*_, *π*_*bj*_ and an individual-specific LD parameter *δ*_*ab,j*_. Thus, to implement this framework, we need a way to generate each of the quantities *π*_*aj*_, *π*_*bj*_, *δ*_*ab,j*_. For example, we could first produce the allele frequencies *π*_*aj*_, *π*_*bj*_ via an admixture model, then draw *δ*_*ab,j*_ from the interval [*D*_min_(*π*_*aj*_, *π*_*bj*_), *D*_max_(*π*_*aj*_, *π*_*bj*_)]. Having obtained IHFs *η*_*ab,j*_, we can then draw haplotypes according to Equation 2, and finally genotypes as linear combinations of haplotypes. We provide details of our particular implementation, used in the statistical power simulations, in Appendix B.

Importantly, individual genotypes simulated under this model will still *marginally* follow the conditional distribution of *X*_*ij*_|*π*_*ij*_ under the coancestry model and they will still be in structural HWE (sHWE). We formalize this property in Proposition A.1. As such, deviations from the null hypothesis for pairs of genotypes drawn through this framework will be due to structural LD, and *not* violations of the sHWE model assumption.

### A.2 Additional theoretical results

We first state a result used to derive the variance of the sLD test statistic 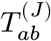 in A test for structural linkage disequilibrium:

#### Lemma A.1.

*If X and Y are a pair of zero-mean, uncorrelated random variables, and further X*^2^ *and Y* ^2^ *are uncorrelated, then the variance of their product is given by*

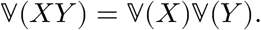

The following result emphasizes that individual genotypes generated through the Multinomial model in Equation 2 follow the conditional distribution of *X*_*ij*_|*π*_*ij*_ under the coancestry model. Note that this result holds for the Multinomial formulation in general. Naturally, it holds for the previously outlined forward-generating probability model of genotypes in structural LD, which offers a specific parameterization of IHFs under this Multinomial framework.

#### Proposition A.1.

*Suppose we form genotypes X*_*aj*_ *and X*_*bj*_ *according to the Multinomial distribution in Equation 2, with X*_*aj*_ = *H*_*ab,j*_ + *H*_*aB,j*_ *and X*_*bj*_ = *H*_*ab,j*_ + *H*_*Ab,j*_. *If the constraints π*_*aj*_ = *η*_*ab,j*_ + *η*_*aB,j*_ *and π*_*bj*_ = *η*_*ab,j*_ + *η*_*Ab,j*_ *are satisfied, then the genotypes will be in structural HWE:*

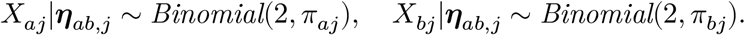

The last result below shows that the null hypothesis of zero conditional covariance between loci *a* and *b* implies the stronger conclusion of *conditional independence* between them under our modeling approach. As such, even though the hypotheses being tested are framed in terms of the conditional covariance between loci *a* and *b*, we can equivalently view the test as one of conditional independence.

#### Proposition A.2.

*Suppose the assumptions of Proposition A*.*1 are satisfied. Then we have that*

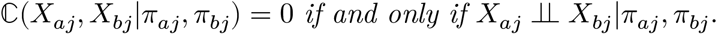

### A.3 Proofs

*Proof of Lemma A*.*1*. We will write

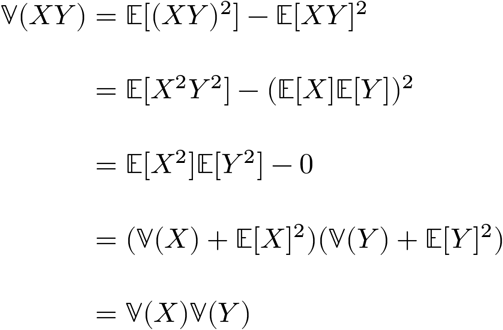

*Proof of Proposition A*.*1*. This result follows from the properties of the Multinomial distribution. Without loss of generality, consider the genotype *X*_*aj*_ = *H*_*ab,j*_ + *H*_*aB,j*_, which is the sum of two disjoint components of a Multinomial vector. Now, let 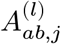 and 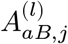 denote indicators for haplotypes *ab* or *aB* being drawn during trial *l ∈* {1, 2} of the Multinomial distribution, respectively. Similarly, let 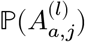 represent the probability of allele *a* appearing in trial *l*, with:

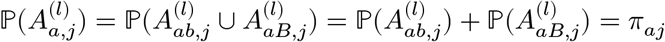

since the events are disjoint. Because the trials (maternal and paternal haplotypes) are independent, 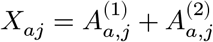 is essentially the sum of two independent Bernoulli random variables each with success probability *π*_*aj*_. Thus,

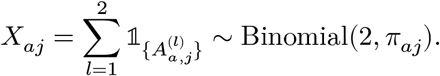

*Proof of Proposition A*.*2*. Since independence trivially implies zero covariance, we will prove the converse. We will use the notation introduced in the preceding proof, writing genotypes as sums of allelic states:

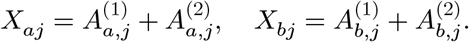

Then we can expand the covariance between genotypes as

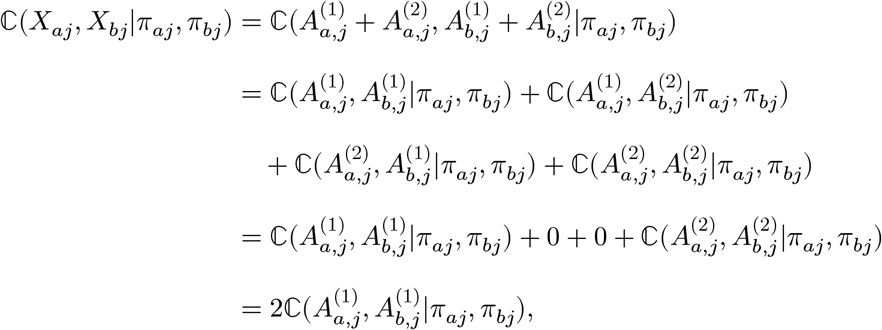

where the third line follows from independence of Multinomial trials and the final line from the symmetry of the maternal and paternal haplotypes. We will focus on the first haplotype without loss of generality. As such, it is clear that

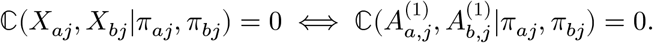

We want to show now that 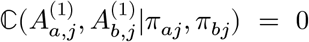 implies conditional independence between *X*_*aj*_ and *X*_*bj*_, using the fact that 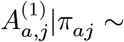 Bernoulli(*π*_*aj*_) and 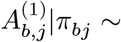Bernoulli(*π*_*bj*_). Thus, we can write the covariance in terms of joint and marginal probabilities:

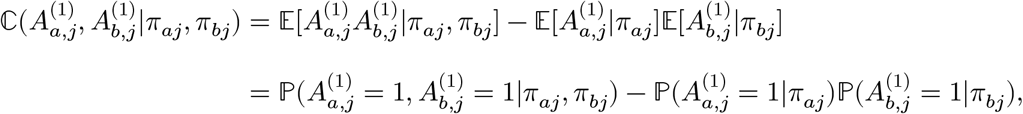

which equals zero iff 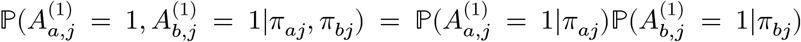 Since the joint probability factors into a product of marginal probabilities, this means that 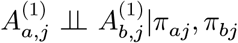 whenever ℂ (*X*_*aj*_, *X*_*bj*_|*π*_*aj*_, *π*_*bj*_) = 0. By symmetry, we also have that 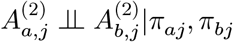 whenever ℂ (*X*_*aj*_, *X*_*bj*_|*π*_*aj*_, *π*_*bj*_) = 0. Because the genotypes 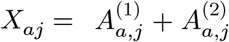 and are 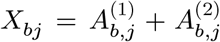 linear combinations of the allelic states, if each pair of allelic states is conditionally independent, the genotypes themselves are likewise conditionally independent.

*Proof of Theorem 2*.*1*. The sequence of random variables 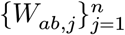 is conditionally independent given 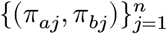 but not necessarily identically distributed. Each variable has finite mean E[*W*_*ab,j*_|*π*_*aj*_, *π*_*bj*_] = 0 and variance

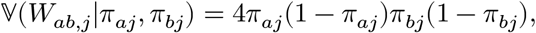

which is uniformly bounded above and below in *j*, with values in the interval [4*ϵ*^2^(1−*ϵ*)^2^, 1/4].

Thus, the variance of the partial sum

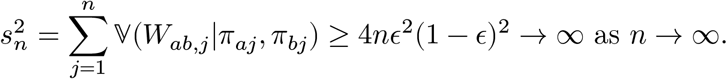

It suffices to show now that the Lindeberg condition [25] for the Central Limit Theorem holds for the sequence of random variables 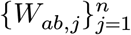, that is, for all *δ* > 0,

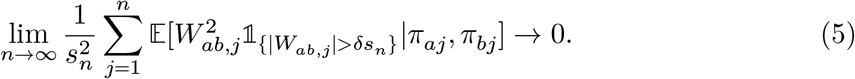

From the uniform bounds on the genotypes and IAFs, it follows that

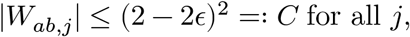

for some finite *C* > 0. Now fix *δ* > 0. Since *s*_*n*_ → ∞, there exists *N*_*δ*_ ∈ ℕ such that for all *n* ≥ *N*_*δ*_, *δs*_*n*_ > *C*. As such, for all *j* and *n* ≥ *N*_*δ*_, we have that 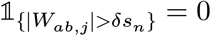 almost surely. Therefore, for all *δ* > 0 and large enough *n*, the term

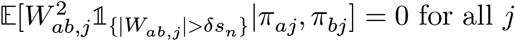

and thus

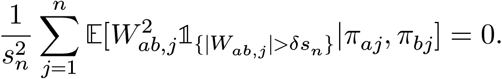

Clearly Equation 5 holds and we conclude that 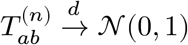 as *n* → ∞.

## B Supplementary Data Analyses

### B.1 Admixture model details

To estimate Π under the admixture model, we use ALStructure [20], which requires Π to be of low rank *K* < *J ≪ I*, where *K* is the number of admixed populations, and *J* and *I* are the numbers of individuals and loci, respectively. ALStructure is a semiparametric approach that projects the genotype matrix **X** onto a factorization Π = **PQ**, with **P** and **Q** denoting *I* × *K* and *K* × *J* matrices of admixed population allele frequencies and admixture proportions, respectively, yielding a consistent estimate 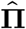 under certain conditions. Related methods, such as ADMIXTURE [32] or fastSTRUCTURE [33], can similarly be used to accomplish this task.

To select the hyperparameter *K*, we apply the sHWE test [22], which evaluates the goodness-of-fit of the conditional distribution of *X*_*ij*_|*π*_*ij*_ in Equation 1 on a locus-by-locus basis across the entire genome. Thus, sHWE offers a procedure for identifying the smallest number of admixed populations *K* such that the vast majority of loci are accurately modeled. While we utilize an admixture model to estimate IAFs in our analyses, we emphasize that our theory and methods are broadly applicable to the more general coancestry model of which admixture is a special case.

### B.2 Simulation study details

Here, we provide technical details regarding the simulations conducted in this study. For Figure 3, we simulate genotypes under an admixture model with *K* = 4 admixed populations for *J* = 800 individuals at *I* = 80000 loci, using the bnpsd library [17]. We draw ancestral allele frequencies from the Uniform(0.05, 0.95) distribution, admixed population allele frequencies via the Balding-Nichols distribution [34], and admixture proportions according to a Gaussian random walk along a one-dimensional geography [16]. We use admixed population inbreeding coefficients of 0.1, 0.2, 0.3 and 0.4. Simulated loci are filtered so that the overall population minor allele frequency (MAF) for each locus is at least 0.01. IAFs are in turn estimated from the simulated genotypes using alstructure [20], under the assumption that the number of admixed populations (*K* = 4) is known.

For Figure 4, we utilize the forward-generating probability model in Appendix A. We first generate IAFs for *J* = 800 individuals at *I* = 200000 loci under an admixture model with *K* = 4 admixed populations. Our simulation parameters and distributions are nearly identical to those we use for Figure 3, only with an increased number of loci *I* and without the MAF-based filtering noted previously. From these IAFs, we draw pairs of genotypes both under the null hypothesis (not in structural LD) and under the alternative (in structural LD). Genotypes *not* in structural LD are jointly independent Binomial draws conditional on their IAFs, in accordance with the coancestry model. For locus pairs in structural LD, we adjust their individual-specific LD parameters *δ*_*ab,j*_ to evaluate the sLD test’s statistical power across varying levels of structural LD.

We now detail the method for calculating the individual-specific LD parameter *δ*_*ab,j*_. As outlined in Appendix A, the feasible values of *δ*_*ab,j*_ are bounded by *D*_min_(*π*_*aj*_, *π*_*bj*_) and *D*_max_(*π*_*aj*_, *π*_*bj*_) – the minimum and maximum possible values of the LD parameter given two allele frequencies *π*_*aj*_ and *π*_*bj*_. Thus, given a pair of IAFs *π*_*aj*_ and *π*_*bj*_ from the admixture model and assuming positive LD, we first compute the theoretical maximum LD parameter *D*_max_(*π*_*aj*_, *π*_*bj*_) as an upper bound for *δ*_*ab,j*_. For locus pairs in structural LD, we code SNPs so that genotypes and IAFs denote the count and frequency of their respective minor alleles. This prevents the maximum positive LD parameter *D*_max_(*π*_*aj*_, *π*_*bj*_) = min{*π*_*aj*_(1 − *π*_*bj*_), (1 − *π*_*aj*_)*π*_*bj*_} from becoming extremely small when the IAFs *π*_*aj*_ and *π*_*bj*_ are highly unequal (i.e., one near 0 and the other near 1). Therefore, we can tune *δ*_*ab,j*_ in a standardized manner without having statistical power confounded by the coding of alleles as either 0 or 1 during the process of generating genotypes.

To vary *δ*_*ab,j*_ in a way that is comparable across individuals who may have different IAFs at loci *a* and *b*, we parameterize it as a function of a shared conditional correlation parameter termed *ρ*_*ab*_ ∈ [−1, 1]. From the Multinomial formulation in Equation 2 (see A test for structural linkage disequilibrium and Appendix A for details), *δ*_*ab,j*_ represents the conditional covariance between loci *a* and *b* within a given haplotype for individual *j*. Using the notation in the proofs of Proposition A.1 and Proposition A.2, we can write the conditional *correlation ρ*_*ab,j*_ between loci *a* and *b* within a given haplotype for individual *j* in terms of the conditional covariance *δ*_*ab,j*_ and IAFs *π*_*aj*_, *π*_*bj*_:

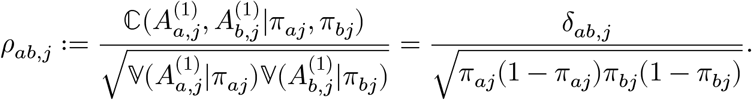

The variance terms follow directly from 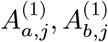 being Bernoulli distributed conditional on *π*_*aj*_, *π*_*bj*_. While we write out moments for the haplotype labeled “1”, this result holds identically for haplotype “2”. Rearranging terms in the equation above yields 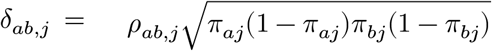. To impose a common level of structural LD for different individuals, we set a target conditional correlation *ρ*_*ab*_ shared across all individuals (i.e., we set *ρ*_*ab,j*_ = *ρ*_*ab*_ for all *j*), while truncating individual-specific LD parameters *δ*_*ab,j*_ that fall outside their feasible range. Thus, for all individuals *j* and non-negative *ρ*_*ab*_, we have

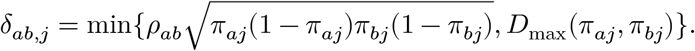

In our simulations, we evaluate values of *ρ*_*ab*_ from 0.1 to 0.5 in increments of 0.05; both empirical FDR and statistical power plateau beyond this range. We compute the theoretical test statistic (using known IAFs) as well as the implemented test statistic, where IAFs are estimated using the entire simulated genotype matrix (including both null and non-null genotype pairs) via `alstructure` [20]. The hypotheses tested consist of 50000 true null hypotheses (genotype pairs not in structural LD) and 2500 true alternative hypotheses (genotype pairs in structural LD). We implement the framework in ref. [30] to control the FDR.

### B.3 Supplementary analyses of human studies

#### B.3.1 Datasets and sampling schemes for locus pairs

All three human studies (TGP, HGDP, HO) were pre-processed and filtered in the same manner as in ref. [22]. We used the BioMart database [35] to annotate SNPs, obtaining their chromosome numbers and locations along a given chromosome. The repository linked in Data Availability contains links to these datasets as well as our implementations.

Our sampling procedures for locus pairs in various settings are as follows. When evaluating interchromosomal LD, we randomly draw 2 × 10^6^ pairs of loci where each locus pair spans two distinct chromosomes. In Figure 8, Figure 9, and Figure 10, we show additional histograms of *p*-values corresponding to interchromosomal locus pairs for the TGP, HGDP, and HO studies, respectively. Each figure contains histograms over a wide range of admixed population counts *K*. For our analysis of LD on the same chromosome (intrachromosomal LD), we draw pairs of loci within a given chromosome. We draw 5000 of such pairs for each chromosome, and do so for all 22 autosomes. In Figure 6, where we plot intrachromosomal LD for each human study, we first transform *p*-values for each locus pair to the − log_10_ scale and bin them according to the base pair distance between their constituent loci. We then plot the mean within each distance window, yielding a smooth curve. We truncate − log_10_(*p*-values) at 16 to account for *p*-values that either equal or approach zero. When investigating the HLA Class II region on chromosome 6, we obtain genomic coordinates for this region from [36], and identify an approximately 1 Mb window within this region.

**Figure 8.**
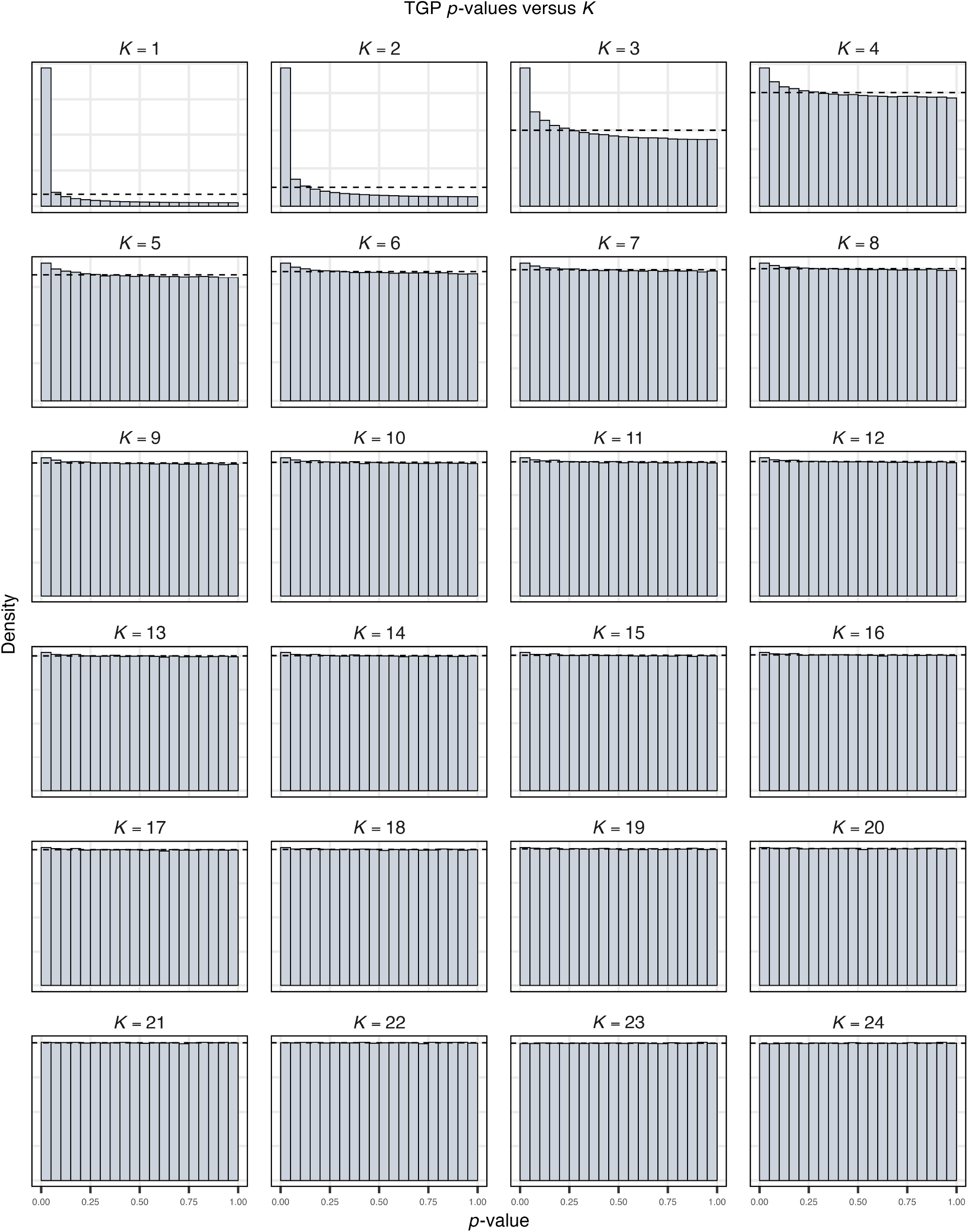
Histograms of interchromosomal locus pair sLD test *p*-values versus the number of admixed populations *K* modeled for the TGP.

**Figure 9.**
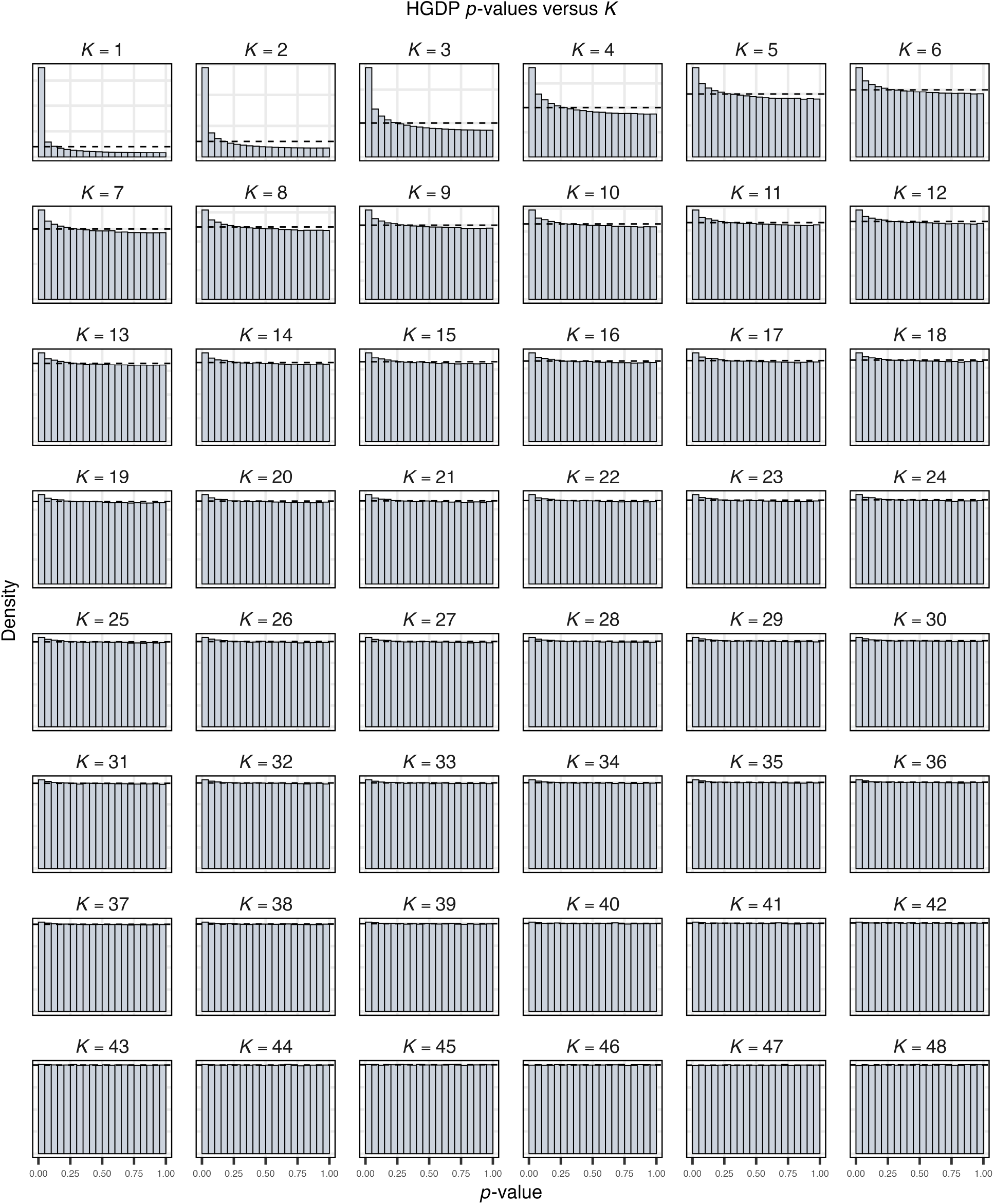
Histograms of interchromosomal locus pair sLD test *p*-values versus the number of admixed populations *K* modeled for the HGDP.

**Figure 10.**
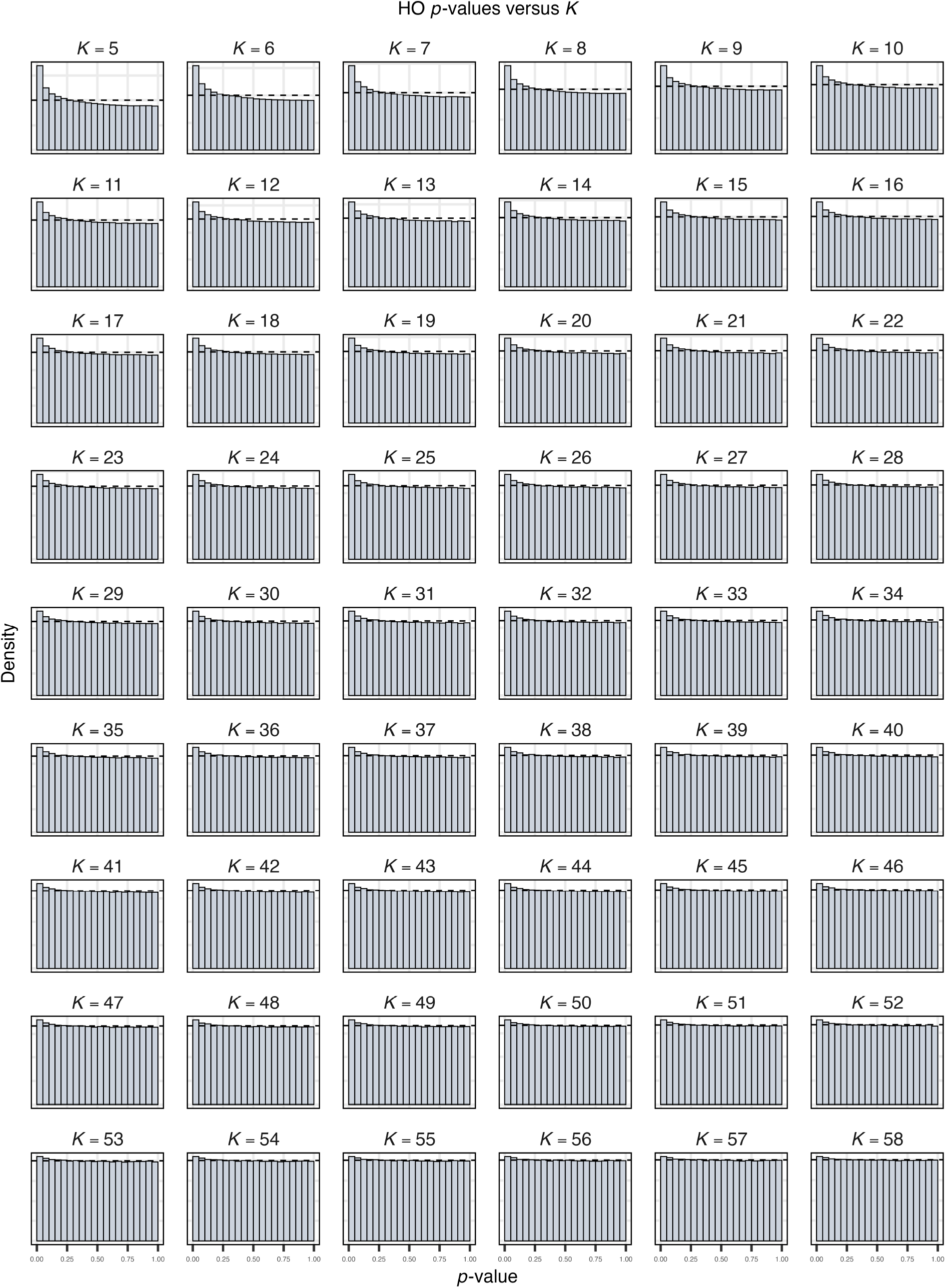
Histograms of interchromosomal locus pair sLD test *p*-values versus the number of admixed populations *K* modeled for the HO study.

#### B.3.2 Evaluating the theoretical Normal(0, 1) null

Here, we evaluate the theoretical Normal(0, 1) null distribution under the setting where the IAFs are estimated by comparing it to an empirical null distribution. Our approach for simulating an empirical null distribution, detailed below, broadly follows the parametric bootstrap outlined in ref. [22]. We analyze the three human studies (TGP, HGDP, HO) to assess whether the Normal(0, 1) approximation is valid in each instance.

We first simulate a null genotype matrix **X**^0^ using IAFs 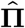 estimated from the TGP, HGDP or HO study. We analyze each of these three studies but do so one at at time, and evaluate a range of admixed population counts *K* for each study. For a given null genotype matrix, we draw its entries jointly independent of one another as 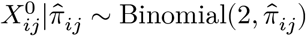. We then re-estimate the IAFs using the null genotypes **X**^0^, yielding a new estimated set of IAFs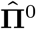. When we estimate 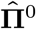, we use the same number of admixed populations *K* as was used to produce the initial estimated IAFs 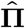. We then compute the implemented sLD test statistic 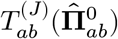 for randomly selected pairs of distinct loci *a* and *b* using the simulated genotypes 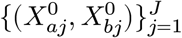 and their respective IAF estimates 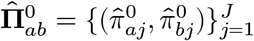. The distribution of these null test statistics constitutes the bootstrap empirical null distribution used to compute *p*-values for observed test statistics. Note that the original algorithm in ref. [22] draws additional null genotype matrices **X**^0^ from 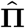 to ensure a numerically stable empirical null. We find that our empirical null is stable and that our results below hold even when using a single null replicate **X**^0^. This is because the number of unique locus pairs we can draw from **X**^0^ forms an empirical distribution whose resolution far exceeds the number of SNPs.

We then assess whether the bootstrap empirical null distribution is approximately Normal(0, 1) distributed. We compare these distributions in − log_10_(*p*-value) space. As such, we compute two-sided *p*-values via the transformation 2(1 −Φ(|*⋅*|)), where Φ(*⋅*) denotes the Normal(0, 1) CDF, and then take the negative log of the *p*-values. Plotting *p*-values on a log axis allows us to better visualize the tails of these distributions. In Figure 11, Figure 12, and Figure 13, we show the concordance between the boostrap empirical null and theoretical Normal(0, 1) distributions. This result holds regardless of the number of admixed populations *K*, demonstrating that the theoretical Normal(0, 1) null is valid irrespective of the population model used. Therefore, there is strong evidence that the theoretical Normal(0, 1) null distribution is appropriate for all three human studies.

**Figure 11.**
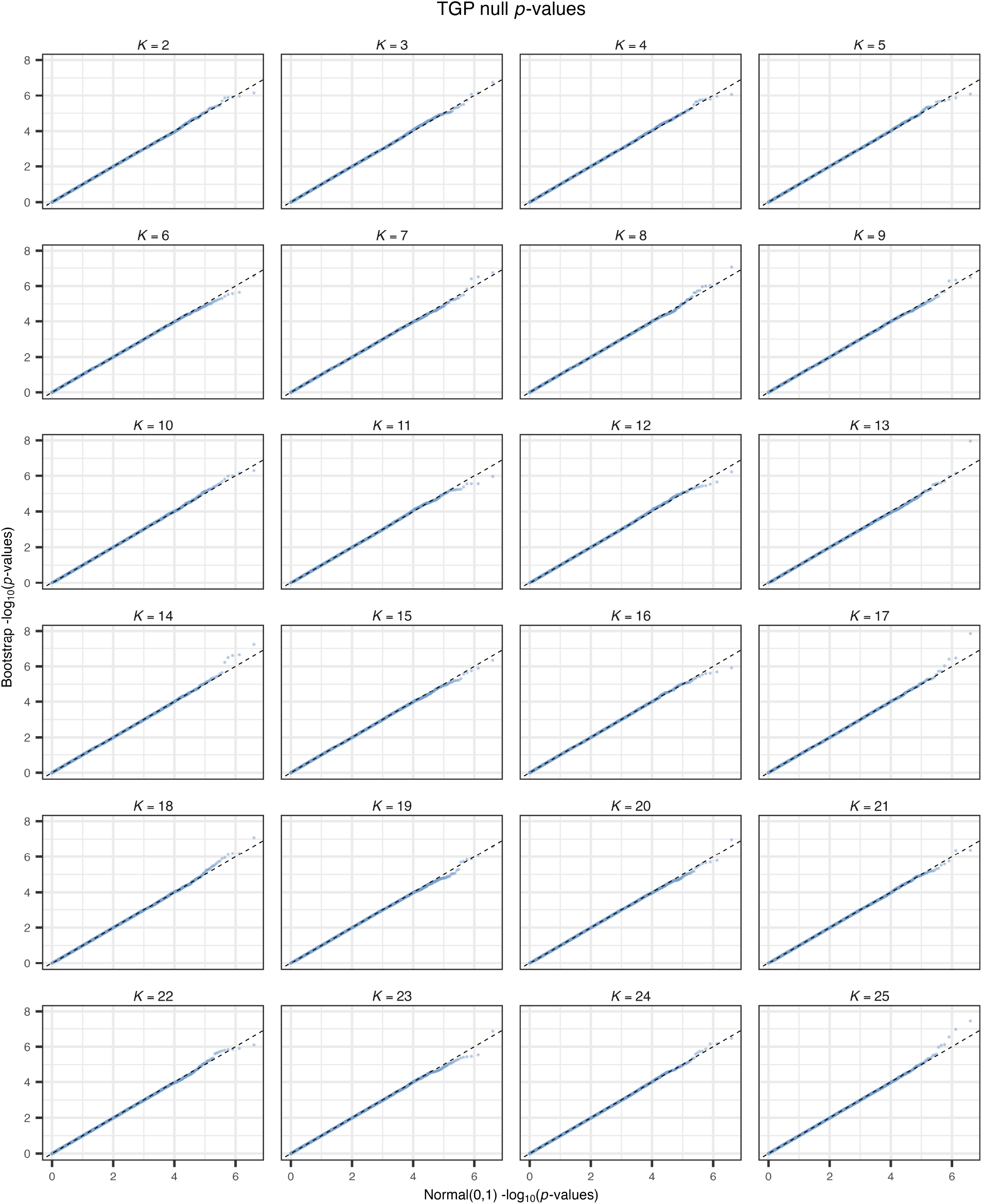
Quantile-quantile plots evaluating the concordance of the theoretical Normal(0, 1) null distribution with bootstrap empirical null distributions for the TGP. We compare the distributions in – log_10_ (*p*-value) space.

**Figure 12.**
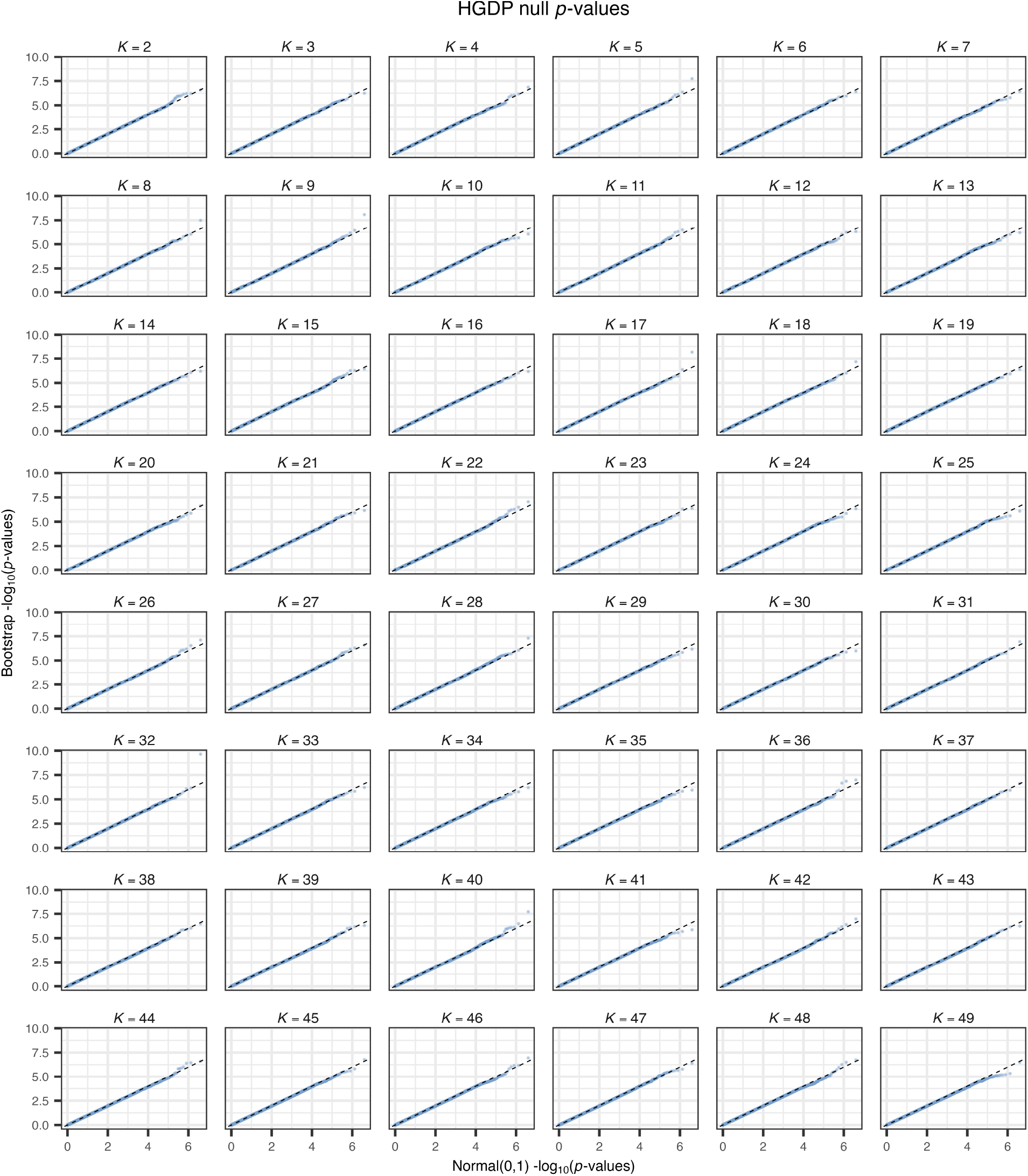
Quantile-quantile plots evaluating the concordance of the theoretical Normal(0, 1) null distribution with bootstrap empirical null distributions for the HGDP. We compare the distributions in − log_10_ (*p*-value) space.

**Figure 13.**
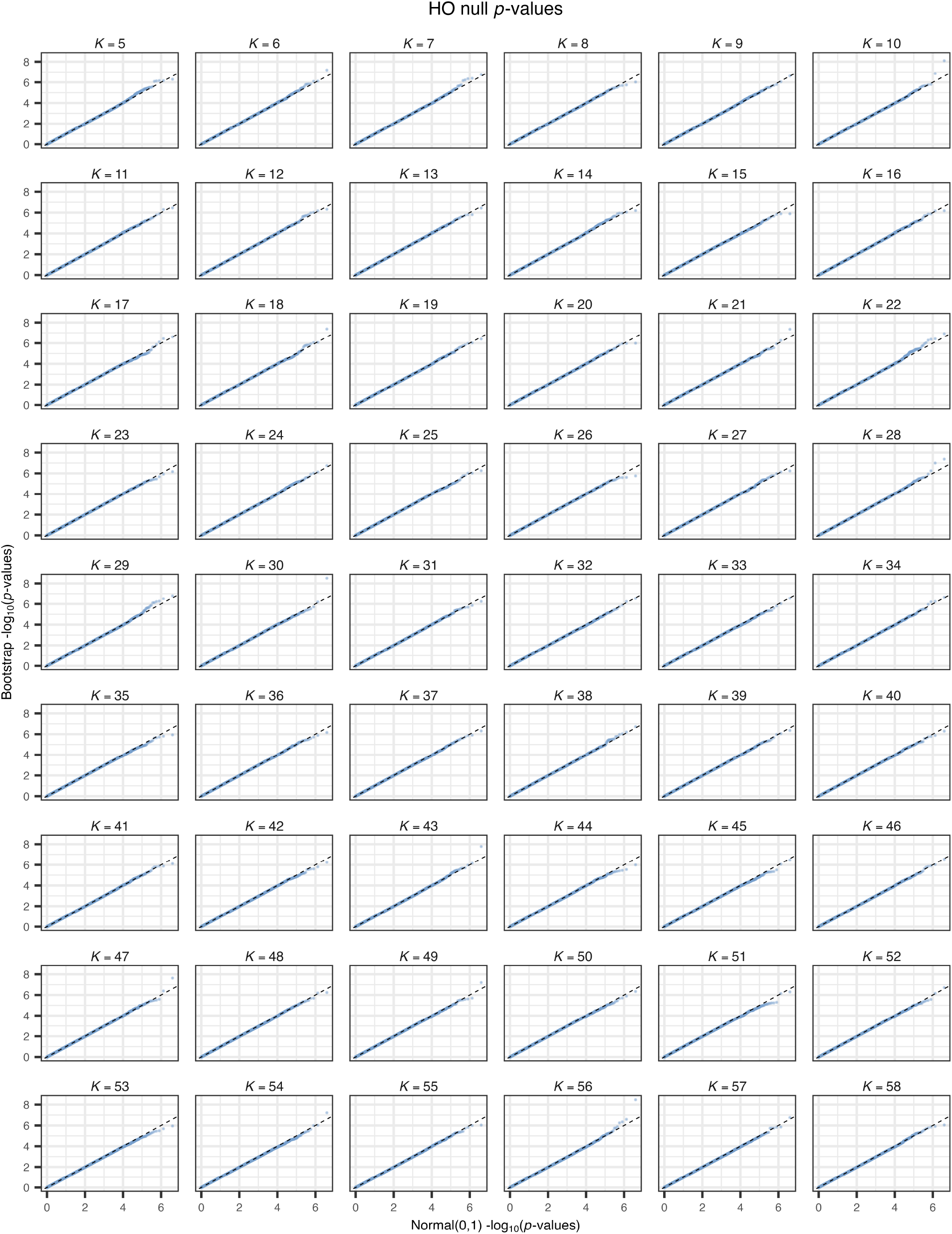
Quantile-quantile plots evaluating the concordance of the theoretical Normal(0, 1) null distribution with bootstrap empirical null distributions for the HO study. We compare the distributions in − log_10_ (*p*-value) space.

## Notes

https://github.com/StoreyLab/sLD-manuscript-analysis

https://github.com/StoreyLab/sLDtest

### Competing Interest Statement

The authors have declared no competing interest.

